# Genetic Control of Radical Crosslinking in a Semi-Synthetic Hydrogel

**DOI:** 10.1101/752436

**Authors:** Austin J. Graham, Christopher M. Dundas, Alexander Hillsley, Dain S. Kasprak, Adrianne M. Rosales, Benjamin K. Keitz

## Abstract

Enhancing materials with the qualities of living systems, including sensing, computation, and adaptation, is an important challenge in designing next-generation technologies. Living materials seek to address this challenge by incorporating live cells as actuating components that control material function. For abiotic materials, this requires new methods that couple genetic and metabolic processes to material properties. Toward this goal, we demonstrate that extracellular electron transfer (EET) from *Shewanella oneidensis* can be leveraged to control radical crosslinking of a methacrylate-functionalized hyaluronic acid hydrogel. Crosslinking rates and hydrogel mechanics, specifically storage modulus, were dependent on a variety of chemical and biological factors, including *S. oneidensis* genotype. Bacteria remained viable and metabolically active in the crosslinked network for a least one week, while cell tracking revealed that EET genes also encode control over hydrogel microstructure. Moreover, construction of an inducible gene circuit allowed transcriptional control of storage modulus and crosslinking rate via the tailored expression of a key electron transfer protein, MtrC. Finally, we quantitatively modeled dependence of hydrogel stiffness on steady-state gene expression, and generalized this result by demonstrating the strong relationship between relative gene expression and material properties. This general mechanism for radical crosslinking provides a foundation for programming the form and function of synthetic materials through genetic control over extracellular electron transfer.

**Significance Statement:** Next-generation materials will require coupling the advantages of engineered and natural systems to solve complex challenges in energy, health, and the environment. Living cells, such as bacteria, naturally possess many of the qualities essential to addressing these challenges, including sensing, computation, and actuation, using their genetic and metabolic machinery. In addition, bacteria are attractive for incorporation into materials due to their durability, ease-of-use, and programmability. Here, we develop a platform for controlling hydrogel properties (e.g., stiffness, crosslinking rate) using extracellular electron transfer from the bacterium *Shewanella oneidensis.* In our system, metabolic electron flux from *S. oneidensis* to a metal catalyst generates radical species that crosslink an acrylate-based macromer to form the gel. This synthetic reaction is under direct control of bacterial genetics and metabolism, which we demonstrate through inducible circuits and quantitative modeling of gene expression and resultant hydrogel properties. Developing methods that capitalize on the programmability of biological systems to control synthetic material properties will enable hybrid material designs with unprecedented functions.

## Introduction

Nature uses hierarchical and genetically-encoded instructions to construct functional materials with specific self-assembly, regulatory, healing, and morphological properties(1). Inspired by such processes, engineered living materials (ELMs) employ the autonomy of living cells to synthesize and control material structures across multiple scales with user-designed functions that are directly coupled to gene expression(2–5). Living materials containing microbes, including biofilms, bacterial cellulose, curli fibers, and synthetic gels loaded with bacteria, are of prominent interest due to their potential application in tissue engineering, 3D printing, soft robotics, metabolic engineering, and living sensors(6–10). Bacteria are particularly attractive as ELM components due to their natural sensing capabilities and programmability. For example, engineered bacteria can act as cellular actuators integrated within the ELM, tailoring its synthesis and function through overexpression, mutagenesis, and gene circuitry.

Not surprisingly, the majority of ELMs rely on materials natively produced by the host organism. For example, several amyloid-based materials have been synthesized by genetically tractable bacteria, such as aggregates of CsgA in *Escherichia coli(11, 12)* and TasA in *Bacillus subtilis(13).* Genetic fusions have allowed these fibrous matrices to bind specific molecules, conduct electricity, perform catalytic reactions, and adhere to complex surfaces(14–17). Apart from amyloids, extracellular polymerization of bacterial cellulose has been engineered using quorum sensing circuits and mutagenesis to create sturdy materials for tissue engineering and sensing applications(18, 19). Despite these advances, significant drawbacks of natural materials include their limited chemical functionality, robustness, homogeneity, and scalability compared to engineered synthetic materials(20). For example, manufactured soft materials such as polymers and hydrogels are easily-functionalized and versatile, facilitating their adoption in diverse environments. However, synthetic materials largely lack the dynamic adaptability and environmental responsiveness found in natural systems. Introducing these qualities to synthetic materials could synergistically enhance ELMs and enable new applications that combine the precision and chemical diversity of engineered materials with the autonomy and evolvability of living cells. However, such designs will require methods for bacteria to control synthetic material properties at the genotypic level. Similarly, robust transcriptional control and quantitative prediction of the relationship between gene expression and material properties are needed for ELMs to approach the design precision of engineered materials.

Toward this goal, we recently developed a cell-controlled radical polymerization reaction using extracellular electron transfer (EET) from the organism *Shewanella oneidensis(21).* In this process, electron flux from native carbon metabolism was redirected to a metal catalyst which controlled a polymerization governed by the atom-transfer radical polymerization (ATRP) mechanism. Importantly, we demonstrated that control over polymer production was directly coupled to cell metabolism and genetically encoded through specific EET proteins. Since ATRP is a versatile platform for soft material synthesis(22), we hypothesized that EET-powered catalysis could be extended to control radical crosslinking in a synthetic hydrogel. While there are numerous examples of incorporating live cells into polymer networks, network properties such as crosslink density, mesh size, degradation, and elastic modulus have generally been designed independent of cell activity. In addition, previous attempts to incorporate cells as live crosslinking agents in synthetic hydrogels have relied on the activity of glucose peroxidase or extracellular functionalization of cells after growth(23–27), which compromise cell viability through the creation of toxic reactive oxygen species or are not under cellular control. Cell-free gelation systems using bacterial lysates have also been explored(28), but removing the living component prevents continued responsiveness. We envisioned that controlling radical crosslinking via EET gene expression would capitalize on the programmability of bacteria and enable the use of stimuli-responsive synthetic biology circuits to control material function.

Here, we demonstrate that EET from *S. oneidensis* can be used to control radical crosslinking of a semi-synthetic methacrylated hyaluronic acid (MeHA) hydrogel (Fig. 1a). First, we show that EET is required for gelation, and that organisms without this metabolic capability (i.e., *E. coli)* are unable to crosslink gels on a comparable time scale or in a controllable manner. Gels did not form unless a constant source of electron flux, radical initiator, and metal catalyst were present. Additionally, the facultative metabolic capability of *S. oneidensis* enabled crosslinking under benchtop conditions without dedicated oxygen removal. Analysis of cell motility and metabolic activity reveled that bacteria remain viable and responsive in the gels for a minimum of one week, and that degree of crosslinking by EET affected cell movement. Next, we found that crosslink density was a strong function of bacterial genetics, as cytochrome knockout strains synthesized gels more slowly and with decreasing stiffness correspondent to the number of removed EET genes. Finally, transcriptional circuits based on controlling the expression of *mtrC* with the LacI repressor enabled tunable crosslinking rates and hydrogel mechanical properties. We found that hydrogel storage modulus fit well to inducible gene expression models, directly linking steady-state gene expression to a quantifiable and macroscopic material property. Overall, our results suggest that transcriptional control over EET can be used to predictably interface the properties of living systems with potentially any material amenable to radical crosslinking.

**Figure 1.**
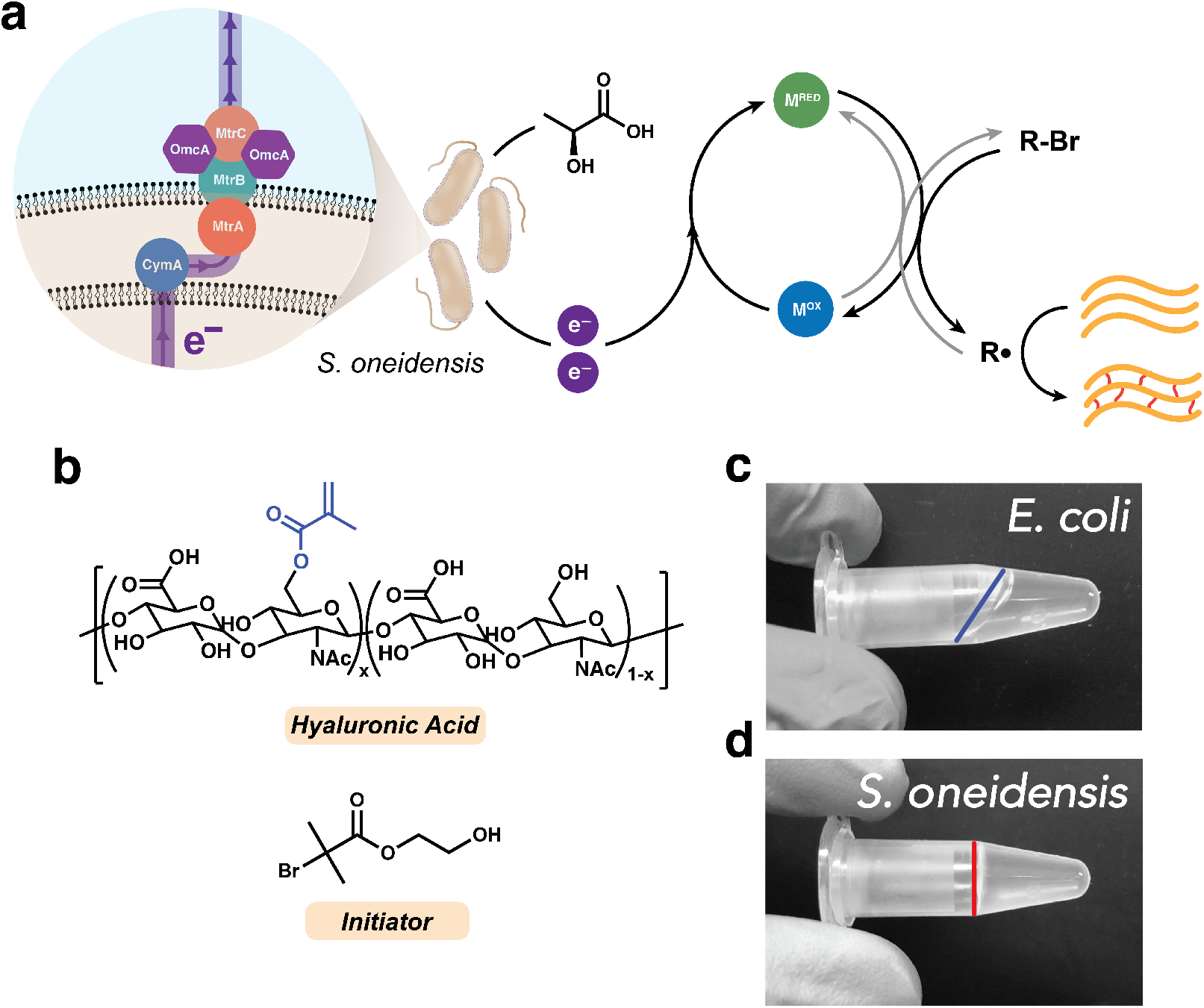
Extracellular electron transfer from *S. oneidensis* controls radical crosslinking of a semi-synthetic hydrogel. (a) The Mtr pathway of *S. oneidensis* transfers metabolic electron flux to a metal catalyst, which generates a radical from a brominated initiator and crosslinks acrylate-based functional groups. (b) Chemical structures of the macromer, methacrylated hyaluronic acid (MeHA), and the radical initiator, 2-hydroxyethyl 2-bromoisobutyrate (HEBIB). (c) Crosslinking reaction mixture inoculated with *E. coli,* which does not possess EET machinery, does not form gels as indicated by liquid flow. Air-liquid interface is highlighted. (d) Crosslinking reaction mixture inoculated with *S. oneidensis* MR-1 forms a solid gel as confirmed by inversion test. Air-liquid interface is highlighted.

## Results

### Extracellular Electron Transfer from Live *S. oneidensis* Controls Aerobic Radical Crosslinking

To initially validate our hypothesis that EET-controlled ATRP could be used to form a crosslinked hydrogel, we first synthesized a 65% methacrylated hyaluronic acid (MeHA) macromer using an established protocol(29) (Fig. S1). Hyaluronic acid is a common naturally-derived biomaterial platform that is attractive for our application due to its biocompatibility and chemical versatility(30). The high density of functional groups was chosen to increase likelihood of successful crosslinking and to minimize the effect of radical scavenging by oxygen. In initial experiments, MeHA was dissolved at 3 wt.% in *Shewanella* Basal Medium (SBM) supplemented with casamino acids (Table S2,3), and the dissolved macromer was mixed with a radical initiator, 2-hydroxyethyl 2-bromoisobutyrate (HEBIB, 500 μM), a copper catalyst with Tris(2-pyridylmethyl)amine ligand (Cu-TPMA, 10 μM), and inoculated with anaerobically pregrown *S. oneidensis* MR-1 cells (OD_600_ = 0.2). Lactate (20 mM) was the electron donor and fumarate (40 mM) was the primary electron acceptor. After mixing, the solution was placed in a humidified anaerobic chamber and was monitored via inversion testing. After 2 h, solutions containing *S. oneidensis* crosslinked to form polymer networks, whereas solutions containing *E. coli* did not (Fig. 1b). Consistent with our previous results(21), these data suggest that electron flux to the metal catalyst from EET-based metabolism is required for radical generation and crosslinking.

Taking advantage of its facultative metabolism, we next tested radical crosslinking of hydrogels by *S. oneidensis* under ambient as opposed to anaerobic conditions. Using the same reagent concentrations as above with aerobically pregrown cells (henceforth, standard conditions), *S. oneidensis* formed crosslinked networks at 30 °C in microcentrifuge tubes without dedicated oxygen removal, as confirmed via inversion test. After these preliminary demonstrations, we more thoroughly investigated the mechanical properties of the gels using shear oscillatory rheology. 50 μL solutions were inoculated and placed between two hydrophobically-treated glass slides with a 0.5 mm silicone spacer, allowed to crosslink, and swollen overnight at room temperature in 1x PBS. Storage and loss moduli were determined from the linear viscoelastic regime as determined by strain and frequency sweeps (Fig. S2), and calculated using a 0.01 to 100 Hz frequency sweep at 0.1% strain. Gels formed in both aerobic and anaerobic environments yielded comparable mechanical properties and were predominantly elastic networks (Fig. S3). Gels prepared by *S. oneidensis* were also mechanically similar to acellular gels crosslinked using UV light and the photoinitiator lithium phenyl-2,4,6-trimethylbenzoylphosphinate (LAP, 500 μM) (Fig. S4). In addition, controls lacking (a) EET-active bacteria, (b) radical initiator, (c) metal catalyst, or (d) methacrylate functional group did not form measurable gels within 2 h (Fig. 2a, Fig. S5). Together, these results demonstrate that EET-crosslinked hydrogels can be synthesized under ambient or anaerobic conditions using electroactive bacteria to form mechanically robust networks that are typical of this macromer.

**Figure 2.**
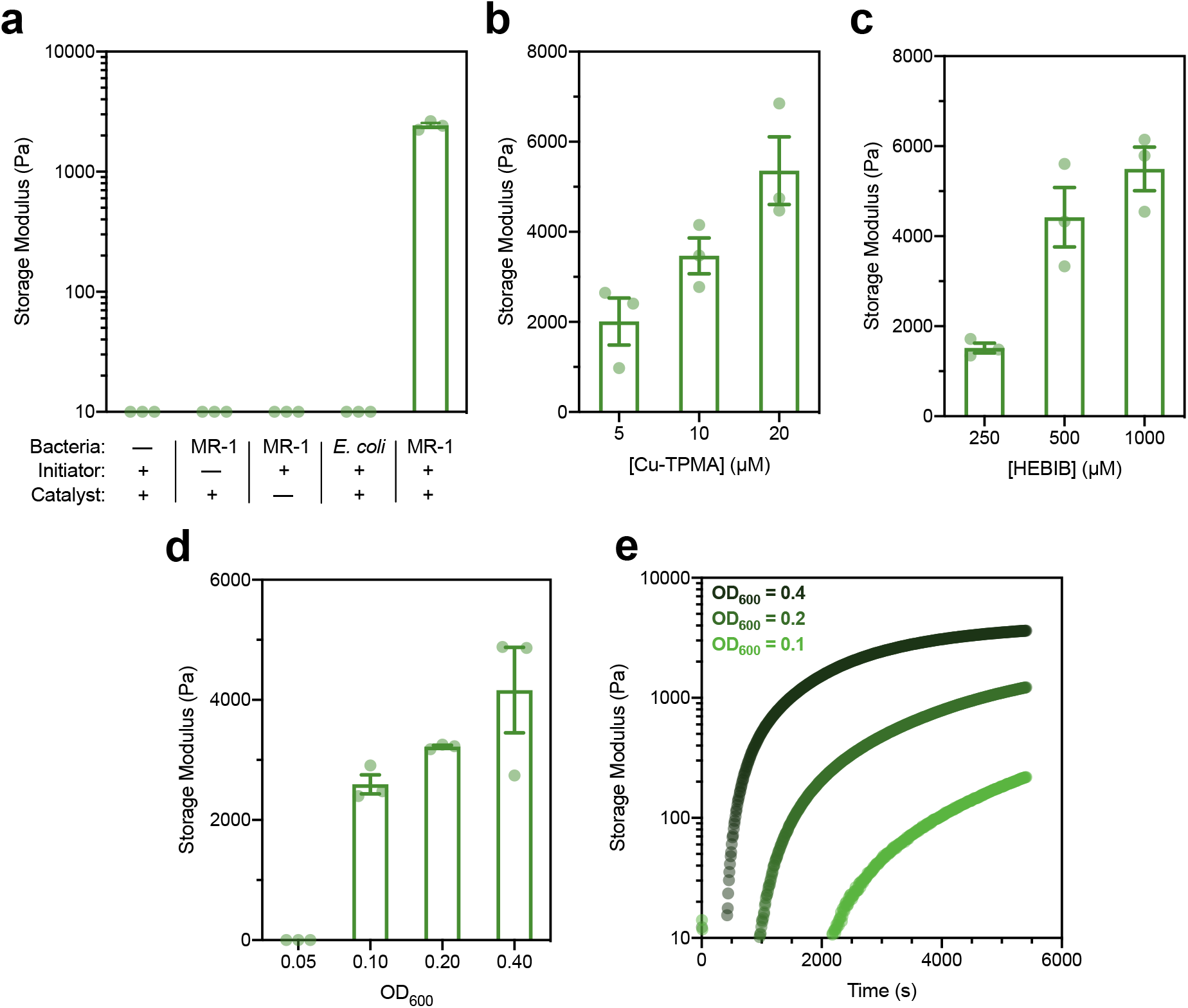
Living hydrogel materials crosslinked by *S. oneidensis* MR-1 can be chemically or biologically tuned. (a) Storage moduli of hydrogels crosslinked with various components missing measured by rheology after 2 hours of crosslinking. Many gels did not form and could not be characterized. (b) Storage moduli of hydrogels crosslinked for 2 hours with various concentrations of Cu-TPMA (catalyst), one-way ANOVA *p* = 0.018. (c) Storage moduli of hydrogels crosslinked for 2 hours with various concentrations of HEBIB (radical initiator), one-way ANOVA *p* = 0.0027. (d) End-point storage moduli at 2 hours and (e) *in situ* rheology of hydrogels crosslinked with various inoculating densities of *S. oneidensis* MR-1 and therefore varying degrees of electron transfer, one-way ANOVA *p* = 0.0002. Gels did not form using OD_600_ = 0.05 in either experiment. (a-d) Data are shown as mean ± SEM, *n* = 3 biological replicates.

The crosslinking kinetics and mechanical properties of polymer networks strongly depend on a variety of chemical factors such as catalyst and ligand identity, rate of initiation, and initiator structure(31). Thus, we next explored the tunable range of hydrogel stiffness after 2 h of crosslinking by altering the concentration and identity of the chemical components in our system (Fig. S6). First, we varied the concentration of the metal catalyst from 5 μM to 20 μM. Greater catalyst concentrations were not considered due to copper-induced transcriptional responses in *Shewanella* at concentrations greater than 20 μM(32). Increasing catalyst concentration correspondingly increased gel modulus (Fig. 2b). Second, we varied the concentration of the initiator over the range of 250 to 1000 μM. As expected, increasing gel stiffness was a function of increasing initiator concentration (Fig. 2c). An additional advantage of ATRP over hydroxyl radicals is the potential for using structurally well-defined radical initiators. Consistent with this expectation, we found that a PEG-based initiator, poly(ethylene glycol) bis(2-bromoisobutyrate) (M_n,avg_ = 700 g/mol), also successfully crosslinked EET-controlled gels at a variety of concentrations (Fig. S7). Overall, EET-controlled hydrogels exhibited a modulus range of about 1-6 kPa for the conditions tested, which is typical of chain-growth crosslinked MeHA hydrogels and within range for a variety of applications, such as biofilm and tissue mimetics(29, 33, 34). These results also indicate that traditional approaches to tuning hydrogel mechanics are still applicable when using EET-controlled crosslinking.

Next, we investigated the role of inoculating *S. oneidensis* cell density, and thus aggregate EET flux, on hydrogel modulus. Below a certain critical inoculum (around OD_600_ = 0.1), the rate of bacterial oxygen consumption was not fast enough to overcome oxygen diffusion and radical quenching. As expected, hydrogels formed with sufficient cell density, and stiffness strongly correlated with OD_600_ (Fig. 2d). Based on these results, we predicted that crosslinking rate would also be coupled to EET and initial cell density. To confirm this, we performed rheological measurements *in situ*, which provided real-time measurement of mechanical properties during crosslinking, at 1 Hz and 0.1% strain. Gels formed at higher initial cell concentrations were not only stronger, but formed more quickly (Fig. 2e). Consistent with end-point experiments, *in situ* rheology measurements also confirmed that a critical concentration of cells was necessary for oxygen depletion. Together, these results demonstrate that cells play a direct role in crosslinking, and overall stiffness and crosslinking rate can be controlled by cell inoculum. They also suggest that genetic and metabolic manipulations to tune EET flux could be used to influence gel mechanics.

### *S. oneidensis* Remain Viable and Metabolically Active in the Polymer Network

For a living material to maintain responsiveness, it is critical that the actuating components (i.e., cells) remain viable and encased in the network. The various components of our system, including the Cu catalyst, initiator, and presence of radicals could affect cell viability. Thus, we assessed cell viability and activity after crosslinking. EET-crosslinked gels formed in standard conditions were swollen overnight in 1x PBS after modulus measurements, and stained using BacLight Live/Dead dyes. Even after mechanical stresses induced by swelling and rheometer measurements, cells maintained approximately 100% viability 5 days after crosslinking (Fig. S9). In addition, cells exposed to crosslinking conditions, but released from the gel surface during swelling, could successfully inoculate new cultures in fresh growth media, indicating viability; these cultures were also able to crosslink new hydrogels with identical properties (Fig. S10).

Since new cultures could be inoculated using cells released during swelling, we next quantified escape or leakage of bacteria from the gels after crosslinking. At the functional group density and crosslink molecular weight of our material, the mesh size of a fully converted gel should be on the order of 10-50 nm(35, 36). Because this is considerably smaller than average bacterial dimensions, there should be minimal cell escape. To test this prediction, crosslinked gels were prepared at standard conditions, swollen in 1 mL of 1x PBS, and the optical density of the surrounding media was measured. An initial, low optical density of cells was detected immediately upon swelling. We hypothesized that this was due to an instantaneous egress of cells on the periphery of the gels and not contained in the network. After washing gels 3x with 1 mL PBS to remove this outer layer of cells, no increase in optical density was detected. Furthermore, colony counting confirmed that escaped cells after 24 h of swelling accounted for < 0.005% of the inoculating density (Fig. S11), suggesting embedded cells do not escape the network in significant numbers.

Continued network adaptation and design of new functions requires an understanding of spatiotemporal cell behavior within the gels during synthesis. Therefore, we next visualized the relationship between genotype, crosslink density, and cell movement during gelation. We constructed an inducible *sfgfp* expression plasmid under the control of the LacI repressor protein and its cognate promoter, P_tac_. Cells were transformed with this vector such that sensing of isopropyl β-D-1-thiogalactopyranoside (IPTG) would induce a fluorescent response indicative of metabolic activity. We developed two reporter strains by transforming both *S. oneidensis* MR-1 and Δ*mtrC*Δ*omcA*Δ*mtrF* (an EET-deficient knockout, described below) with this construct. Strains were grown overnight in 1000 μM IPTG, washed, and inoculated into standard gelation mixtures. The solution was then pipetted onto a glass slide and sealed under a coverslip, such that crosslinking occurred in the sealed layer. Bacterial movement was monitored by time-lapse imaging using GFP fluorescence. Cells were uniformly dispersed within the network throughout gelation. For both *S. oneidensis* MR-1 and Δ*mtrC*Δ*omcA*Δ*mtrF*, a significant degree of bacterial motion was visible upon inoculation, both by convective flow of the reaction mixture and by flagella-based swimming(37). Minutes after inoculation, cell movement and bulk fluid motion was arrested in the *S. oneidensis* MR-1 sample as crosslinking proceeded (Movie S1). Contrastingly, movement both from flow and swimming were still perceptible after 2 h in the Δ*mtrC*Δ*omcA*Δ*mtrF* sample, indicating that minimal crosslinking occurred (Movie S2). Cell movement was quantified using TrackMate in Fiji 1.0(38), which revealed that average cell displacement over 5 seconds was significantly greater for the knockout strain at both 0 and 2 h (Fig. 3a, Fig. S12). Movement was not significantly different between the two strains in non-functionalized hyaluronic acid solution, suggesting the observed motility differences were due to crosslinking, even at early times immediately following inoculation (Fig. S13, Movies S3-4). These results confirm that cells become trapped in the polymer network as it forms, and suggest that bacterial genotype encodes control over bulk and microscopic properties such as crosslink density and mesh size, affecting flow, diffusion, and cell movement within the material.

**Figure 3.**
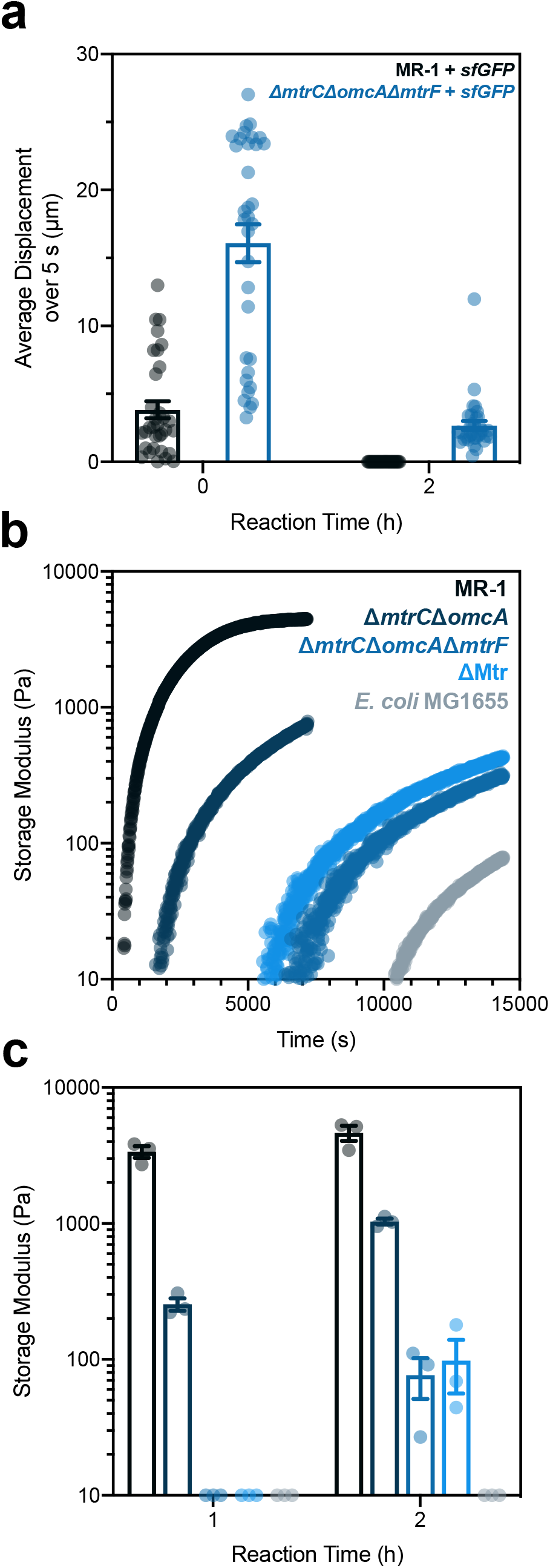
Crosslink density, synthesis rate, and cell motility within living hydrogels are governed by *S. oneidensis* genetics and EET machinery. (a) Average cell displacement within the gels measured by microscopy over 5 second time-lapses at both 0 and 2 hours into crosslinking. Displacement was quantified using TrackMate in Fiji 1.0. Student *t*-test *p* < 0.0001 between strains at *t* = 0 and 2 hours. (b) *In situ* and (c) end-point rheology measurements of hydrogels crosslinked by *S. oneidensis* strains with various EET genes knocked out. *E. coli* was included as an EET-deficient control. Student t-test *p* < 0.0001 for MR-1 compared to other strains at *t* = 1 and 2 hours. Data are shown as mean ± SEM, (a) *n* = 33 tracked cells or (c) *n* = 3 biological replicates.

Although the cells remained viable for days in the crosslinked gels, we wished to assess their continued sensing and metabolic capabilities over long periods after crosslinking. Gels synthesized at standard conditions with sfGFP-expressing *S. oneidensis* MR-1 were swollen in 1x PBS for varying lengths of time after crosslinking, then induced for 24 h with 1000 μM IPTG. Significant fluorescence was detected by microscopy in induced samples up to 1 week after crosslinking (Fig. S14), but was not detectable in uninduced samples. Together, these results indicate that the bacteria remain viable, trapped, and maintain transcriptional and translational capabilities for extended periods after crosslinking.

### Bacterial Genetics Govern Crosslink Density

Understanding the genetic link between EET and crosslinking is critical for biologically controlling hydrogel structure and function. Toward this goal, we employed various EET knockout strains in crosslinking reactions. The Mtr pathway is a primary source of EET flux in anaerobic *S. oneidensis* metabolism (Fig. 1a). Outer membrane cytochromes MtrC and OmcA are terminal reductases of the Mtr pathway, and responsible for direct transfer of electrons onto metal species such as Fe and Cu(21, 39). MtrF is a homologue to MtrC and can similarly reduce a variety of metals(40). We employed three different cytochrome knockout strains in assessing the role of EET in crosslinking: Δ*mtrC*Δ*omcA*, Δ*mtrC*Δ*omcA*Δ*mtrF*, and ΔMtr. The knockout ΔMtr refers to a strain with a large number of EET genes knocked out that should provide minimal electron flux to the catalyst (Table S1)(39, 41). *E. coli* MG1655 was also included as an EET-deficient control. We measured *in situ* crosslinking kinetics using these strains, and compared crosslinking rates and density. Both crosslinking rate and hydrogel storage modulus strongly corresponded with bacterial genotype, where decreasing number of EET genes led to decreased crosslinking rates and weaker moduli (Fig. 3b). Although MtrC is the primary terminal reductase for many metal substrates, our results show that MtrF exhibits compensatory reduction of Cu in the *ΔmtrCΔomcA* knockout compared to the Δ*mtrC*Δ*omcA*Δ*mtrF* knockout. The strong similarity between gels formed by the Δ*mtrC*Δ*omcA*Δ*mtrF* and ΔMtr knockouts further demonstrates that outer membrane cytochromes are primarily responsible for electron transfer to the Cu catalyst and subsequent crosslinking activity. The minimal, delayed crosslinking activity of *E. coli* suggests that background radical generation or non-specific Cu reduction can produce weak gels at extended times. In separate experiments, we corroborated these *in situ* results using end-point, swollen gel measurements after 1 and 2 h of crosslinking (Fig. 3c, Fig. S15). *S. oneidensis* MR-1 and Δ*mtrC*Δ*omcA* formed gels the fastest and were measurable at 1 h, whereas the other strains did not form measureable gels by this time. Measurable networks were formed by Δ*mtrC*Δ*omcA*Δ*mtrF* and ΔMtr at 2 h, but were significantly weaker than gels formed by the strains containing more EET machinery. Overall, these results show that bacterial genotype directly governs gel modulus and suggests that material properties can be controlled through more sophisticated regulation of EET.

### Transcriptional Regulation of Extracellular Electron Transfer Yields Tunable Crosslinking Activity

For ELMs to emulate the adaptability of biological materials, the actuating components should continually sense and respond to their environment. Environmental stimuli should then induce a transcriptional response and impart control over material properties. Toward this goal, we constructed an inducible *mtrC* expression plasmid using the same genetic circuit outlined before, but replacing *sfgfp* with *mtrC* (Fig. S16-17). We transformed the Δ*mtrC*Δ*omcA*Δ*mtrF* strain with this plasmid, such that IPTG would sequentially activate *mtrC* expression, electron transfer, and crosslinking activity. Upstream of the *mtrC* gene, a computationally-predicted weak synthetic ribosome binding site was employed to optimize control over EET and minimize leaky expression(42). SDS-PAGE and heme staining of total protein from induced and uninduced cell lysates validated inducible MtrC protein production after overnight growth in IPTG-containing medium. High molecular weight bands corresponding to the size of the MtrCAB complex were observed in induced Δ*mtrC*Δ*omc*Δ*mtrF* samples and a wild-type control, but not in uninduced and empty vector Δ*mtrC*Δ*omcA*Δ*mtrF* samples (Fig. S18). Functional steady-state expression of *mtrC* in response to IPTG was further validated by measuring Fe^3+^ reduction with the ferrozine assay. After 2 h of reduction, Fe^2+^ concentration increased with the presence of inducing molecule, indicating functional MtrC activity and no leaky EET response over an uninduced control (Fig. S19). Next, we verified tailored crosslinking activity in response to varying transcriptional activation. *In situ* gelation kinetics were assessed after overnight growth in media containing a range of inducing molecule concentrations. Crosslinking activity was a strong function of IPTG concentration, spanning orders of magnitude in storage modulus (Fig. 4b). Crosslinking kinetics also corresponded to inducer presence, indicating that both synthesis rate and final material modulus can be customized through differential steady-state gene expression. Both an induced empty vector control and a complemented strain with no IPTG did not form measurable gels in 2 h. Thus, transcriptional regulation over EET gene expression in response to an environmental signal imparts programmable control over hydrogel stiffness.

**Figure 4.**
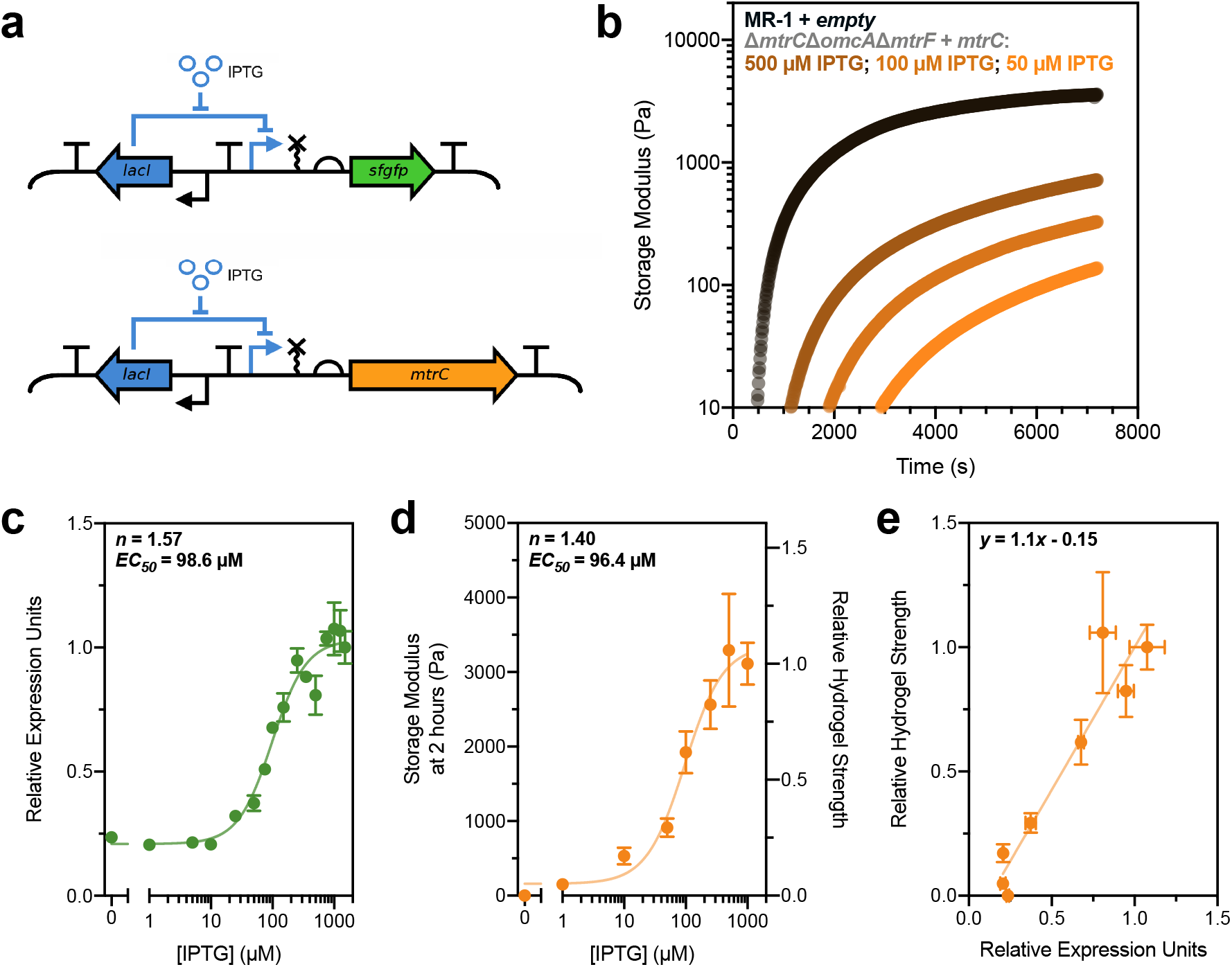
Modeling gene expression allows quantitative prediction of living hydrogel properties. (a) Genetic circuits utilized in this study placed either *sfgfp* or *mtrC* under inducible control of IPTG via the LacI repressor and P_tac_ promoter. (b) *In situ* rheology of hydrogels crosslinked by *S. oneidensis* with steady-state induced *mtrC* levels at various IPTG concentrations. Δ*mtrC*Δ*omcA*Δ*mtrF* + empty and Δ*mtrC*Δ*omcA*Δ*mtrF* + *mtrC* with 0 μM IPTG were also tested, but did not form gels on the time scale shown. (c) Hill function analysis of sfGFP fluorescence (denoted as relative expression units by normalization to fluorescence at maximum induction) as a function of IPTG concentration. (d) Hill function analysis of hydrogel storage modulus after 2 hours of crosslinking as a function of IPTG concentration. Right y-axis is storage modulus normalized to average modulus at maximum induction. (e) Normalized hydrogel stiffness plotted as a function of relative expression units for corresponding IPTG concentrations, fit to a line (*R*^2^ = 0.80). (c-e) Data are shown as mean ± SEM, *n* = 3 biological replicates.

### Modeling Gene Expression Enables Predictable Material Properties

Due to successful transcriptional regulation of *mtrC*, we hypothesized that a material property such as storage modulus could be predicted from inducible gene expression models. Since the *sfgfp* and *mtrC* circuits have identical transcriptional regulation, we tested whether both fit to activating Hill function models. First, we measured the response function of the *sfgfp* circuit in *S. oneidensis* MR-1 by inducing overnight cultures in a variety of IPTG concentrations. Steady-state fluorescence was quantified using a plate reader, and normalized to optical density (Fig. 4c). As expected, relative expression (i.e., normalized fluorescence) was a strong function of IPTG concentration, and fit well to a Hill function with a hillslope of *n* = 1.57 and a half-maximal effective concentration of *EC*_50_ = 98.6 μM (Table S5). These results indicate that our circuit generates a predictable transcriptional response. Next, end-point gel measurements were used to examine storage modulus as a function of steady-state cytochrome (MtrC) expression. Gels were crosslinked for 2 h since our *in situ* results indicated this time would provide sufficient differentiation between induced cultures at varying IPTG concentrations. Specifically, Δ*mtrC*Δ*omcA*Δ*mtrF* complemented with LacI-regulated *mtrC* was grown overnight in a variety of IPTG concentrations and allowed to react for 2 h at standard gelation conditions. We found that hydrogel storage moduli were also under strong transcriptional control, similar to *sfgfp*, and could be modeled using a Hill function with *n* = 1.40 and *EC*_50_ = 96.4 μM (Fig. 4d). As the sfGFP signal is effectively a measure of the transcriptional rate at different IPTG concentrations(43), the similarity between fitted constants for *sfgfp* expression and hydrogel stiffness suggests a model where transcriptionally-controlled MtrC levels predictably control hydrogel properties. To further visualize this relationship, we plotted normalized storage modulus as a function of relative expression units for each corresponding IPTG concentration and observed a linear correlation (Fig. 4e). The 1:1 relationship between steady-state gene expression and hydrogel properties is corroborated by the approximate unity of the slope. Together, these results demonstrate that EET gene expression can be modulated to control ELM properties (e.g., gel stiffness), and that fluorescence-parameterized models for existing and new genetic circuits may be adapted to design, predict, and control more complex macroscopic material outputs.

## Discussion

We showed that *S. oneidensis* can genetically control radical crosslinking in a semisynthetic hydrogel via electron transfer to a redox-active polymerization catalyst. Similar to other crosslinking chemistries, storage modulus was dependent on catalyst and initiator concentrations as well as initial cell density. A significant advantage of our system is that it is theoretically amenable to any substrate that can undergo radical crosslinking and support microbial life. We used methacrylate-functionalized hyaluronic acid, but other semi-synthetic materials based on functionalized alginate, collagen, and cellulose, as well as completely synthetic substrates, such as PEG, should show similar behavior. In addition to flexibility in macromer structure, our design also allows for a variety of well-defined ATRP initiators to be used as crosslinking agents. For example, we showed two traditional ATRP initiators, HEBIB and bis-brominated PEG, could both form crosslinked hyaluronic acid hydrogels. Similar to other radical crosslinking methodologies, bacteria-controlled crosslinking is also compatible with various biochemical modifications including the installation of integrin recognition motifs (e.g., RGD), orthogonal crosslinking chemistries (e.g., Michael addition), and other common polymer engineering paradigms. The chemical flexibility and general compatibility with a variety of polymer network scaffolds should facilitate the use of our platform in tissue engineering, 3D printing, soft robotics, and drug delivery.

In contrast to other biologically-driven radical crosslinking methods, most notably hydroxyl radicals generated from glucose oxidase, EET-controlled crosslinking did not negatively impact cell health. Cells remained viable at least one week following gelation and transformed cells could express *sfgfp* in response to an external stimulus. We also observed genotypic changes in cell motility and convective flow as a result of EET-dependent crosslinking, implying genetic control over gel microstructure. Overall, our design avoids cell viability concerns associated with other radical crosslinking methodologies and could enable synthetic platforms for studying biofilm formation(44), trapping(45), or functionalizing cells(46).

We found that crosslinking activity and overall hydrogel stiffness were governed by EET cytochrome expression. Specifically, *S. oneidensis* MR-1 with wild-type EET pathways generated stiff gels within an hour while negative controls containing *E. coli* MG1655, which lacks EET machinery, did not form gels on comparable timescales. At longer time scales (ca. 3-4 hours), EET-knockout strains and *E. coli* showed some crosslinking activity. This background radical generation could be caused by non-specific copper reduction (e.g., release or secretion of cytosolic reducing agents) or spurious radical activation. To further reduce background crosslinking, decreasing catalyst and/or initiator concentration, lowering cell density, or changing the identity of chemical components could all potentially be tuned. Overall, the strong link between *S. oneidensis* genetics and crosslinking rate and density lays the foundation for developing more sophisticated EET-based regulation of material properties.

Using *in situ* and end-point rheology measurements, we showed that hydrogel crosslinking is directly linked to *mtrC* expression levels, which has a number of implications for adaptable and dynamic materials. Interestingly, placing *mtrC* under the control of the LacI repressor generated a hydrogel stiffness response function that is characteristic of inducible gene expression. This response function mirrored one generated from *sfgfp* expression in the same construct, indicating robust transcriptional control over both gene expression and material properties. More importantly, our results suggest that previously characterized genetic circuits, including genetic logic gates, designed to express fluorescent reporters could be readily adapted to control *mtrC* expression and gel stiffness(47). Although the ultimate stiffness of the hydrogel will depend on the specific crosslinking chemistry, our observation of canonical Hill function responses demonstrates that changes in hydrogel stiffness governed by transcriptional regulation can be partially predicted. Overall, our results suggest that a variety of transcriptional circuits could be extended to control the macroscopic properties of synthetic materials in a predictable and programmable manner. Additionally, robust genetic control over crosslinking should complement other stimuli-responsive hydrogel designs, including integration of biochemical signals(48–50), actuators(51, 52), and complex geometric designs(53).

The gel stiffness response function was measured after 2 h of gelation since *in situ* measurements indicated this would be sufficient time to distinguish between differentially induced cells. Because crosslinking is a dynamic process, the hydrogel response function also varies as a function of time, even at steady-state expression levels of MtrC. For example, at early times (< 1 h), it is exaggerated since measurable gels do not form at low induction levels (Fig. S20). At longer timescales, the response function begins to collapse as background polymerization starts to compete with EET-driven crosslinking. In addition to understanding how EET influences crosslinking dynamics, many applications leveraging genetic control over material properties will likely require real-time transcriptional responses. Understanding how cells optimize transient gene expression to control dynamic outputs such as cell motility, morphogenesis, biofilm structure, or extracellular matrix construction are ongoing challenges in developmental and systems biology(54). Similarly, we are currently investigating how to coordinate transient gene expression to the polymerization kinetics in our system(55). Overall, continued optimization of MtrC (or other EET protein) expression and material chemistry should allow for actuation of material changes over timescales similar to transcription and translation, as well as predictive models that relate gene expression to material function.

Overall, we found that extracellular electron transfer from *S. oneidensis* could power a radical polymerization catalyst and form a semi-synthetic hydrogel composed of functionalized hyaluronic acid. A variety of chemical and biological factors controlled crosslinking kinetics and the resulting storage moduli of the gels, demonstrating a tunable and adaptable platform. Most importantly, we found that robust transcriptional control over *mtrC* expression and metabolic electron flux enabled precise and predictable control over hydrogel mechanical properties. While cells are frequently incorporated into polymer networks, our platform allows for a variety of network properties including crosslink density, mesh size, degradation, diffusion, and elastic modulus to be controlled through cellular metabolism and gene expression. In summary, our results provide a powerful foundation to program adaptive and responsive behavior into the vast functional space of synthetic materials through the conduit of biological electron transfer.

## Supporting information

Graham Hydrogel Supplementary Information

## Acknowledgements

*S. oneidensis* knockouts were a generous gift from Prof. Jeffrey Gralnick (U. Minnesota). A.J.G. was supported through a National Science Foundation Graduate Research Fellowship (Program Award No. DGE-1610403). This research was supported by the Welch Foundation (Grants F-1929), and by the National Science Foundation through the Center for Dynamics and Control of Materials: an NSF Materials Research Science and Engineering Center under DMR-1720595. The authors acknowledge use of shared research facilities supported in part by the Texas Materials Institute, the Center for Dynamics and Control of Materials: an NSF MRSEC (DMR-1720595), and the NSF National Nanotechnology Coordinated Infrastructure (ECCS-1542159). A.M.R. gratefully acknowledges a Career Award at the Scientific Interface (#1015895) from the Burroughs Wellcome Fund. We gratefully acknowledge the use of facilities within the core microscopy lab of the Institute for Cellular and Molecular Biology, University of Texas at Austin. NMR spectra were collected on a Bruker Avance III 500 funded by the NIH (Award 1 S10 OD021508-01) and a Bruker Avance III HD 400 funded by the NSF (Award CHE 1626211).

## Author Contributions

A.J.G., A.M.R., and B.K.K. conceived the project; A.J.G., C.M.D., and D.S.K. performed experiments; C.M.D. designed and built the *sfgfp* and *mtrC* expression plasmids; A.H. assisted with MeHA synthesis and rheology; A.J.G., C.M.D., A.M.R., and B.K.K. analyzed the results; A.J.G., C.M.D., A.M.R., and B.K.K. wrote the manuscript.

## Materials and Methods

### Chemicals and Reagents

Sodium hyaluronate (72 kDa, Lifecore Biomedical), methacrylic anhydride (Sigma-Aldrich, 94%), copper(II) bromide (CuBr_2_, Sigma-Aldrich, 99%), tris(2-pyridylmethyl)amine (TPMA, Sigma-Aldrich, 98%), 2-hydroxyethyl 2-bromoisobutyrate (HEBIB, Sigma-Aldrich, 95%), poly(ethylene glycol) bis(2-bromoisobutyrate) (PEGBBIB, M_n,avg_ = 700 g/mol, Sigma-Aldrich, PDI ≤ 1.1) sodium DL-lactate (NaC_3_H_5_O_3_, TCI, 60% in water), sodium fumarate (Na2C_4_H_2_O_4_, VWR, 98%), HEPES buffer solution (C_8_H_18_N_2_O_4_S, VWR, 1 M in water, pH = 7.3), potassium phosphate dibasic (K_2_HPO_4_, Sigma-Aldrich), potassium phosphate monobasic (KH_2_PO_4_, Sigma-Aldrich), sodium chlrodie (NaCl, VWR), ammonium sulfate ((NH_4_)_2_SO_4_, Fisher Scientific), magnesium (II) sulfate heptahydrate (MgSO_4_·7H_2_O, VWR), trace mineral supplement (ATCC), casamino acids (VWR), silicone oil (Alfa Aesar), isopropyl β-D-1-thiogalactopyranoside (IPTG, Teknova), kanamycin sulfate (C_18_H_38_N_4_O_15_S, Growcells), nail polish (Electron Microscopy Sciences), BacLight Live/Dead Stain (Invitrogen), deuterium oxide (D_2_O, Sigma-Aldrich, 99.9%), hydrogen peroxide solution (H_2_O_2_, Sigma-Aldrich, 30% in H_2_O), and 3,3’,5,5’-Tetramethylbenzidine (TMBZ, Alfa Aesar, 98%) were used as received. All media components were autoclaved or sterilized using 0. 2 μm PES filters.

### Methacrylated Hyaluronic Acid Synthesis and Purification

MeHA was functionalized using methacrylic anhydride according to an established protocol(29). Briefly, ~72 kDa HA macromer (1.5 g, 3.81 mmol, 1.0 eq) was dissolved at 1 wt.% in DI water (150 mL), cooled on ice, and adjusted to pH = 8.5 using 5 N NaOH. The pH was maintained between 7.5 – 8.5 using NaOH while methacrylic anhydride (8.44 mL, 56.7 mmol, 14.9 eq) was added in 750 μL aliquots every ~5 minutes. Once all the methacrylic anhydride was added, the pH was maintained between 7.5 – 8.5 for 4 hours, then the reaction stoppered and stirred overnight at room temperature. The reaction solution was dialyzed using 6 – 8 kDa dialysis tubing in DI water while stirring for two weeks. The mixture was then frozen and lyophilized. Methacrylate functionalization was quantified by ^1^H-NMR spectroscopy and determined to be ~65% from integration of the vinyl group relative to the HA backbone (Fig. S1). Functionalized MeHA solution was passed through an alumina column immediately prior to crosslinking reactions.

### Bacteria Strains and Culture

Bacterial strains and plasmids are listed in Table S1 Cultures were prepared as follows: bacterial stocks stored in 20% glycerol at −80 °C were streaked onto LB agar plates (for wild-type and knockout strains) or LB agar with 25 μg/mL kanamycin (for plasmid-harboring strains) and grown overnight at 30 °C for *Shewanella,* 37 °C for *E. coli.* Single colonies were isolated and inoculated into *Shewanella* Basal Medium (SBM) supplemented with 100 mM HEPES, 0.5% trace mineral supplement, 0.5% casamino acids, and 20 mM 60% sodium lactate (2.85 μL/mL) as the electron donor. Aerobic cultures were pregrown in plastic 15 mL culture tubes at 30 °C and 250 rpm shaking. Anaerobic cultures were pregrown using the same procedure outlined above, but in degassed medium in a humidified anaerobic chamber and supplemented with 40 mM sodium fumarate (40 μL/mL of a 1 M stock) as the electron acceptor. Cultures were washed after pregrowth using SBM supplemented with 0.5% casamino acids (degassed for anaerobic cultures). OD_600_ was measured using a NanoDrop 2000C spectrophotometer and normalized to 2.0 for 10-fold dilution into gel mixtures unless otherwise noted.

### MeHA Hydrogel Crosslinking using *S. oneidensis*

CuBr_2_, TPMA, and HEBIB stock solutions were prepared according to previously established protocol(21). For three 50 μL hydrogel discs that were analyzed by rheology, a reaction mixture was prepared as follows: MeHA was dissolved at 3.76 wt.% in SBM with 0.5% casamino acids and aliquoted into an autoclaved microfuge tube (119.2 μL). 400 μM Cu-TPMA (3.75 μL), 69 mM HEBIB (1.09 μL), 60% sodium lactate (0.428 μL), and 1 M sodium fumarate (6 μL) were added to the MeHA solution and mixed. Final concentrations in solution were 3 wt.% MeHA, 10 μM Cu-TPMA, 500 μM HEBIB, 20 mM lactate, and 40 mM fumarate. This solution was distributed into three autoclaved microfuge tubes of 45 μL aliquots to which 5 μL of OD_600_ normalized cells were added. The gel solutions were mixed and added to hydrophobically-treated glass slides with a 0.5 mm silicone spacer separating the two glass layers. Slides were sealed with a binder clip and allowed to react at 30 °C for two hours unless otherwise noted. Hydrogels were removed from the slides using a razor blade and placed into 3 mL baths of 1x PBS overnight to swell to equilibrium. Hydrogels analyzed by *in situ* rheology were prepared using the same mixture outlined above, but inoculated with cells and immediately placed on the rheometer for analysis.

### Rheological Analysis

End-point rheological analysis: swollen hydrogels prepared as outlined above were analyzed by oscillatory shear rheology using a TA Instruments Discovery HR-2 Rheometer with an 8 mm parallel plate geometry. Hydrogels were loaded onto a Peltier plate and excised to 8 mm diameter using a biopsy punch. The geometry gap was then lowered until the measured axial force was above 0.02 N (usually between 500 – 800 μm, depending on the crosslink density and swelling ratio). Storage and loss moduli were measured using frequency sweeps from 0.01 to 100 Hz at a constant strain of 0.1%. Moduli for a single gel were quantified by averaging the linear viscoelastic region of each frequency sweep.

*In situ* rheological analysis: hydrogels measured by *in situ* oscillatory shear rheology were prepared using the mixtures and rheometer outlined above. Immediately after inoculating reaction mixtures with cells, 80 μL of mix was loaded onto the Peltier plate, which was maintained at 30 °C. A 20 mm parallel plate geometry was lowered onto the solution while spinning such that the mixture coated the entire geometry surface and filled the gap (~350 μm gap size). The edges of the geometry and gap were then coated with silicone oil to prevent evaporative losses. *In situ* crosslinking was monitored using 1 Hz oscillation and 0.1% strain over variable lengths of time (1.5 – 8 hours).

### Microscopy and Cell Tracking

All microscopy was performed using a Nikon Ti2 Eclipse inverted fluorescence microscope. Cells assessed for viability by microscopy were crosslinked using standard conditions and the resulting gels swollen in 1x PBS at room temperature for varying lengths of time. The gels were then incubated in the dark in the BacLight Live/Dead stain mix (1.5 μL/mL Syto9, 2.5 μL/mL propidium iodide in 0.85% NaCl solution) for 30 minutes. Stained gels were then washed by pipetting 3x in 1 mL PBS to remove unbound dye. Gels were loaded onto glass microscope slides and a no. 1 coverslip placed on top. The gel thickness prevented using nail polish to seal the sides, but evaporative losses were not noticeable over the course of the experiment. Fluorescence for each stain (green for Syto9, red for propidium iodide) was measured using GFP and Texas Red excitation/emission filter cubes on a Nikon Ti2 Eclipse, as outlined previously(21). To assess metabolic activity, gels were crosslinked with sfgfp-harboring strains and allowed to swell in 1x PBS for varying lengths of time. sfGFP fluorescence was assessed before induction to ensure there was no detectable background fluorescence. Gels were then incubated in 1000 μM IPTG in PBS for 24 hours and monitored by fluorescence using the GFP channel.

Cells tracked by microscopy during crosslinking were prepared with reaction mixtures as outlined above. Upon cell inoculation, the crosslinking mixture was loaded onto glass slides, covered, and sealed with nail polish. The slides were loaded onto the microscope and cell movement monitored using the Time-lapse function in NIS-Elements. Images were taken every 1 s or 5 s with 100 or 300 ms exposure time using the GFP channel. Time-lapse images were edited and quantified using TrackMate in Fiji 1.0. Images were first background subtracted and equally brightened by thresholding. The top 50 highest quality cells were selected, as determined by the TrackMate user interface, and tracked over 5 or 10 s. The highest quality tracks, as determined by the software, were used to quantify average total displacement over the time-lapse. The number of tracks used to calculate the average was the maximum number of tracks for the image with the fewest tracks at each time point.

### Plasmid Construction

DNA sequences and plasmid maps for each genetic part and plasmid used in this study are given in the Supplementary Information. All plasmids were assembled via Golden Gate cloning using enzymes and buffers from New England Biolabs. In addition to T4 Ligase, Golden Gate reactions contained either SapI for *mtrC* plasmid assembly or BsaI for *sfgfp* and empty plasmid assembly. The pCD backbone was assembled by PCR amplifying (Phusion, New England Biolabs) regions of pSR58.6 (B0015 and T0 terminators, ColE1 origin of replication), pTKEI-tLOV (kanamycin resistance), and pAL-rfp_(RP4 origin of transfer). To construct the gene expression unit (insulating terminators, RiboJ ribozyme, and *lacI* regulation unit), a gBlock was synthesized (Integrated DNA Technologies) and used in Golden Gate cloning. *sfgfp* and *mtrC* were PCR amplified from pSR58.6 and purified *Shewanella oneidensis* MR-1 genomic DNA, respectively, with ribosome binding sites and Golden Gate restriction enzyme sites added via oligonucleotide primers. Generally, 10 μL Golden Gate reactions were set up that contained 10 fmol of pCD plasmid backbone and 40 fmol of each gBlock and/or PCR insert (as necessary). In a thermocycler, Golden Gate reactions were cycled 25 times: 90 s at 37 °C followed by 3 minutes at 16 °C. After the 25 cycles, reactions were incubated at 37 °C for 5 minutes, 80 °C for 10 minutes, and then held at 4 °C.

Golden Gate reactions were used to directly transform freshly prepared electrocompetent *S. oneidensis* strains(56). To prepare electrocompetent *S. oneidensis,* 5 mL of overnight *S. oneidensis* growth in LB medium at 30 °C was washed 3 times with sterile 10% glycerol at room temperature and concentrated to ~300 μL. 2 μL of Golden Gate reaction was mixed with 30 μL of concentrated electrocompetent *S. oneidensis,* transferred to a 1 mm electroporation cuvette, and electroporated at 1250 V. To recover electroporated cells, 250 μL of LB was immediately added post-electroporation and cells were incubated/shaken at 30 °C and 250 rpm. After 2 h of recovery, 100 μL of cell suspension was plated onto LB agar plates containing 25 μg mL^-1^ kanamycin sulfate and incubated overnight at 30 °C to obtain single colonies (generally 5-100 colonies observered for 1-3 part assemblies). Single colonies were used to inoculate LB liquid medium containing 25 μg mL^-1^ kanamycin sulfate and incubated/shaken overnight at 30 °C and 250 rpm. These cultures were used to generate 22.5% glycerol stocks, which were stored at −80 °C, and harvest assembled plasmid for Sanger sequencing (DNA Sequencing Facilities, University of Texas at Austin).

### Verification of MtrC Inducible Expression and Functional Activity

Heme staining was performed by adapting previously described methods (57). 5 mL of uninduced (0 μM IPTG) and induced (1000 μM IPTG) *S. oneidensis* strains were anaerobically cultured overnight in SBM containing 20 mM lactate and 40 mM fumarate. The total culture was washed once in 1x PBS, concentrated to 500 μL, and lysed by sonication. The cell lysate was centrifuged for 10 minutes at 10,000 rcf, and the supernatant was transferred to a separate tube. The pellet was resuspended in 100 μL 1x PBS, and the total protein concentration of both lysate fractions was determined by Bradford assay. 10 μg of protein from the supernatant and pellet were loaded into each well of a 12% Bis-Tris SDS-PAGE gel and run for ca. 120 minutes at 110 V. The gel was stained in a 3:7 mixture of 6.3 mM TMBZ in methanol:0.25 M sodium acetate (pH 5.0) for 2 h in the dark. Heme-containing protein bands were visualized upon addition of 30 mM hydrogen peroxide for 30 min.

Ferrozine assay(58) was performed to determine functional activity of MtrC-expressing strains. Strains were anaerobically grown overnight in IPTG-free SBM containing 20 mM lactate and 40 mM fumarate. Within in anaerobic chamber, cells were diluted 100-fold into 96-well plate wells filled SBM containing 20 mM lactate, 5 mM Fe(III)-citrate, and 1 mg mL^-1^ ferrozine (final total volume of 250 μL per well). Each well contained either 0 or 750 μM IPTG and Fe(II) standards were also added to the plate. Immediately after addition of cells and Fe(III), the plate was sealed with optically transparent sealing film and a plate cover lined with silicone grease. The plate was then removed from the anaerobic chamber and statically incubated at 30 °C. Absorbance at 562 nm was periodically measured using a BMG LABTECH CLARIOstar plate reader.

### Quantification and Modeling of Inducible Constructs

The Δ*mtrC*Δ*omcA*Δ*mtrF* + *sfgfp* and + *mtrC* strains were anaerobically pregrown overnight using the same conditions outlined above, with the addition of 25 μg mL^-1^ kanamycin and varying IPTG concentrations. Cells were anaerobically washed 3x in degassed SBM with 0.05% casamino acids, normalized to OD_600_ = 2.0, and diluted 10-fold into reaction mixtures (also containing kanamycin and IPTG) prepared in ambient conditions. Prior to measuring sfGFP fluorescence, protein translation was arrested by supplementing a 100 μL aliquot of cell suspension with kanamycin sulfate to a final concentration of 2 mg mL^-1^. Subsequently, this suspension was shaken aerobically for 1 h at 30 °C to allow for sfGFP maturation. sfGFP fluorescence (488/530 nm) and cell suspension absorbance (600 nm) was measured using a BMG LABTECH CLARIOstar plate reader to yield fluorescence absorbance^-1^ for each sample. For each sample, the background fluorescence absorbance^-1^ from an empty vector (pCD8) control was subtracted. The background subtracted values were then normalized to the average fluorescence absorbance^-1^ value at maximum induction (1500 μM IPTG) to give relative expression units. Crosslinking strains complemented with *mtrC* were allowed to form gels for two hours, and the gels allowed to swell overnight in 1x PBS at room temperature. Gels were then analyzed by oscillatory shear rheology as outlined above. A nonlinear fitting algorithm in GraphPad Prism 8.0 was used to fit inducible gene expression and hydrogel storage modulus to the following activating Hill function: 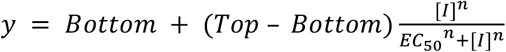. Normalized hydrogel storage modulus and relative expression units were plotted for values at corresponding IPTG concentrations and modeled using a linear fit. Further details on modeling can be found in Note S1. Fitting parameters and “goodness of fit” can be found in Table S5.

### Statistical Analysis

Unless otherwise noted, data are reported as mean ± SEM of *n* = 3 biological replicates. Significance was calculated in GraphPad Prism 8.0 using either a two-tailed unpaired student t-test or a one-way ANOVA (*α* = 0.05).

### Data deposition

Experimental data supporting the findings in this study are publicly available through the Texas Data Repository (doi: XXX).

