## Supplementary material for "Genetic Control of Radical Crosslinking in a Semi-Synthetic Hydrogel": Graham Hydrogel Supplementary Information

**Table S1.** Bacterial strains and plasmids used in this study.

| Strain or plasmid | Description/Genotype | Reference or source |
|---|---|---|
| ***S. oneidensis* Strains** | | |
| MR-1 | MR-1 (ATCC700550), wild-type strain | American-Type Culture Collection |
| JG749 | Lacks outer membrane cytochromes, MtrC and OmcA; Δ*mtrC*Δ*omcA* | [1] |
| JG596 | Lacks out membrane cytochromes MtrC, OmcA, and MtrF; Δ*mtrC*Δ*omcA*Δ*mtrF* | [1] |
| JG1194 | Lacks numerous proteins responsible for EET, including outer membrane cytochromes, β-barrel cytochromes, and periplasmic electron carriers; ΔMtr | [2] |
| MR-1 + pCD7sfGFP | Wild-type with a LacI repressed *sfGFP* circuit (Fig. S16-17) | This work |
| JG596 + pCD7sfGFP | JG596 with a LacI repressed *sfGFP* circuit (Fig. S16-17) | This work |
| MR-1 + pCD8 | Wild-type with an empty vector on the LacI repressed circuit | This work |
| JG596 + pCD8 | JG596 with an empty vector on the LacI repressed circuit (Fig. S17) | This work |
| JG596 + pCD24r1 | JG596 with a LacI repressed *mtrC* circuit (Fig. S16-17) | This work |
| ***E. coli* Strains** | | |
| MG1655 | Wild-type strain | Lydia Contreras, U. of Texas at Austin |
| **Plasmids** | | |
| pCD7sfGFP | See Fig. S16-17 | This work |
| pCD8 | See Fig. S17 | This work |
| pCD24r1 | See Fig. S16-17 | This work |

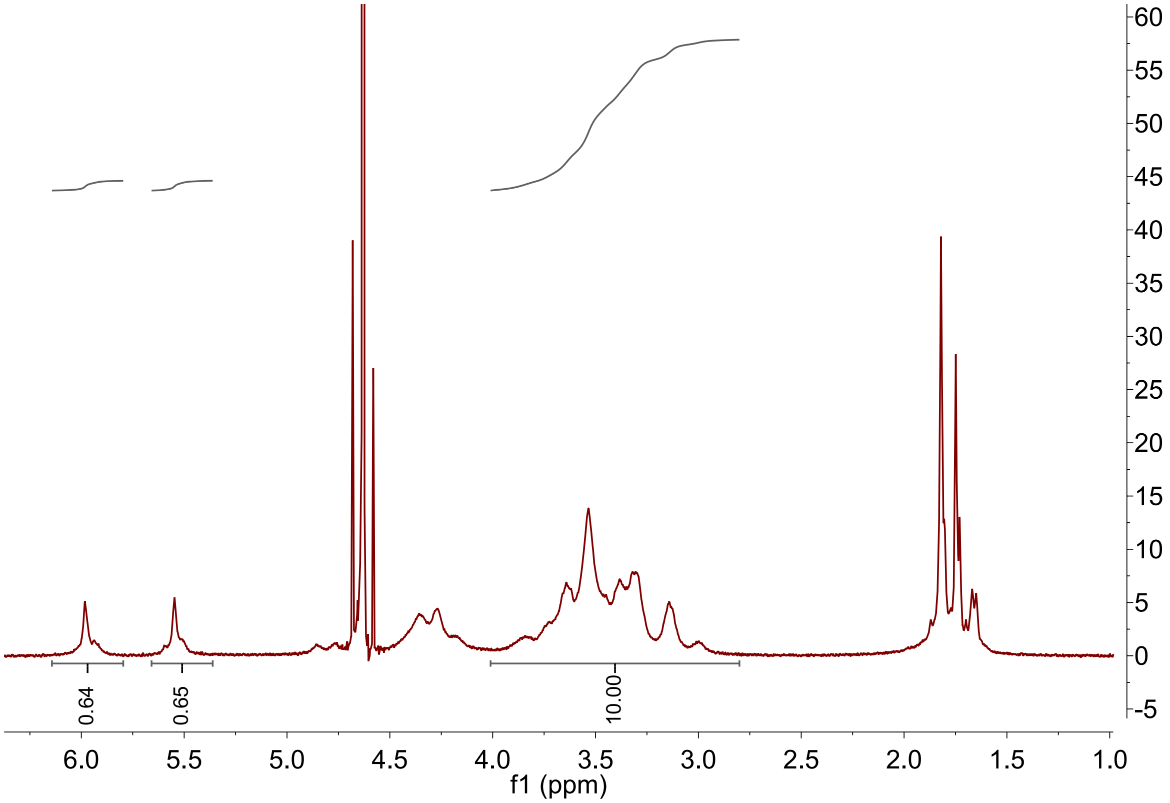


**Figure S1.** $^1$H-NMR spectrum of 65% methacrylated hyaluronic acid macromer dissolved in $D_2O$ after synthesis, dialysis, and lyophilization.

**Table S2.** Ingredients in *Shewanella* Basal Medium. Growth media was supplemented with casamino acids and Wolfe's mineral solution, whereas crosslinking media was only supplemented with casamino acids.

| Ingredient | Quantity for 1L of 1x SBM |
|---|---|
| $K_2HPO_4$ | 225 mg |
| $KH_2PO_4$ | 225 mg |
| NaCl | 460 mg |
| $(NH_4)_2SO_4$ | 1.703 mL of 1 M stock |
| $MgSO_4 \cdot 7H_2O$ | 0.475 mL of 1 M stock |
| HEPES | 100 mL of 1 M stock |
| Casamino acids | 5 mL of 10% stock |
| Wolfe's mineral solution | 5 mL of 200x stock |
| $ddH_2O$ | Up to 1 L, adjust to pH = 7.2 |

**Table S3**. Ingredients in Wolfe's mineral solution, adapted from ATCC recipe.

| Reagent | Quantity in 1L of 200X Stock |
|---|---|
| EDTA | 0.5 g (2.69 mL of 0.5 M stock) |
| $MgSO_4$•$7H_2O$ | 3.0 g |
| $MnSO_4$•$H_2O$ | 0.5 g |
| NaCl | 1.0 g |
| $FeSO_4$•$7H_2O$ | 0.1 g |
| $Co(NO_3)_2$•$6H_2O$ | 0.1 g |
| $CaCl_2$ | 0.9 mL from 1 M stock |
| $ZnSO_4$•$7H_2O$ | 0.1 g |
| $CuSO_4$•$5H_2O$ | 10 mg |
| $AlK(SO_4)_2$ | 10 mg |
| $H_3BO_3$ | 10 mg |
| $Na_2MoO4$•$2H_2O$ | 10 mg |
| $Na_2SeO_3$ | 1 mg |
| $Na_2WO_4$•$2H_2O$ | 10 mg |
| $NiCl_2$•$6H_2O$ | 20 mg |
| $ddH_2O$ | 1 L |

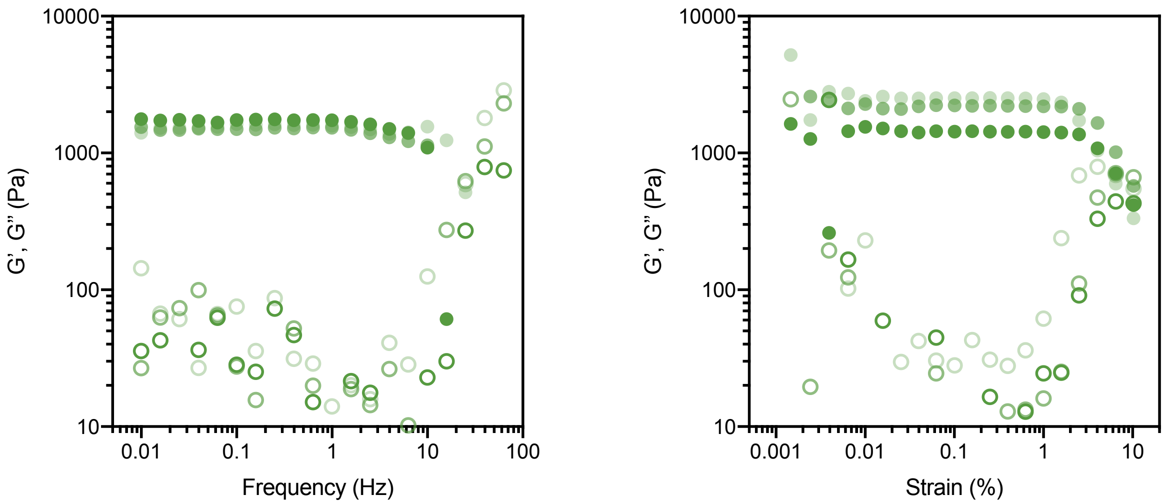


**Figure S2.** Strain and frequency sweeps of MeHA hydrogels crosslinked by *S. oneidensis* MR-1 at standard conditions ($OD_{600}$ = 0.2 cells, 3 wt.% MeHA65, 10 μM Cu-TPMA, 500 μM HEBIB, 20 mM lactate, 40 mM fumarate) for $n$ = 3 biological replicates. Storage modulus calculations for all subsequent gels were made using frequency sweeps at 0.1% strain, which is within the linear viscoelastic regime.

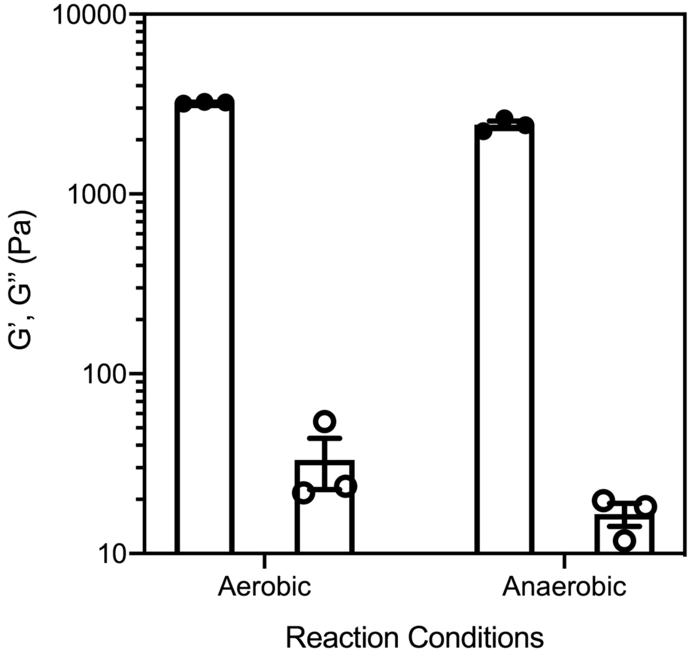


**Figure S3.** G' (filled circles) and G'' (open circles) of hydrogels crosslinked by *S. oneidensis* MR-1 at standard conditions in either aerobic or anaerobic environments. Data shown are mean ± SEM for $n$ = 3 bioloigcal replicates. MeHA65 networks are predominately elastic in nature, as demonstrated by significantly larger G' than G''.

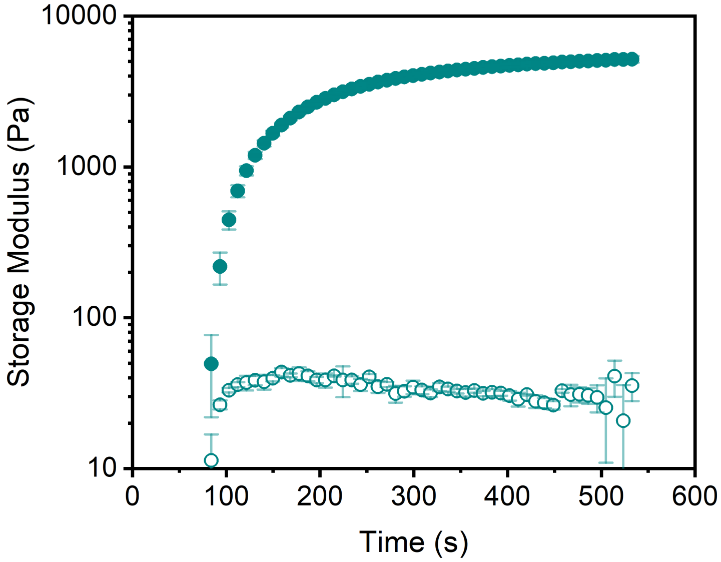


**Figure S4.** *In situ* rheology showing G' (filled circles) and G'' (open circles) of MeHA65 gels synthesized using 500 μM lithium phenyl-2,4,6-trimethylbenzoylphosphinate (LAP) as a photoinitiator with 365 nm light at 10 mW/cm$^2$ laser intensity. Data are reported as the mean ± SD of $n$ = 3 gels.

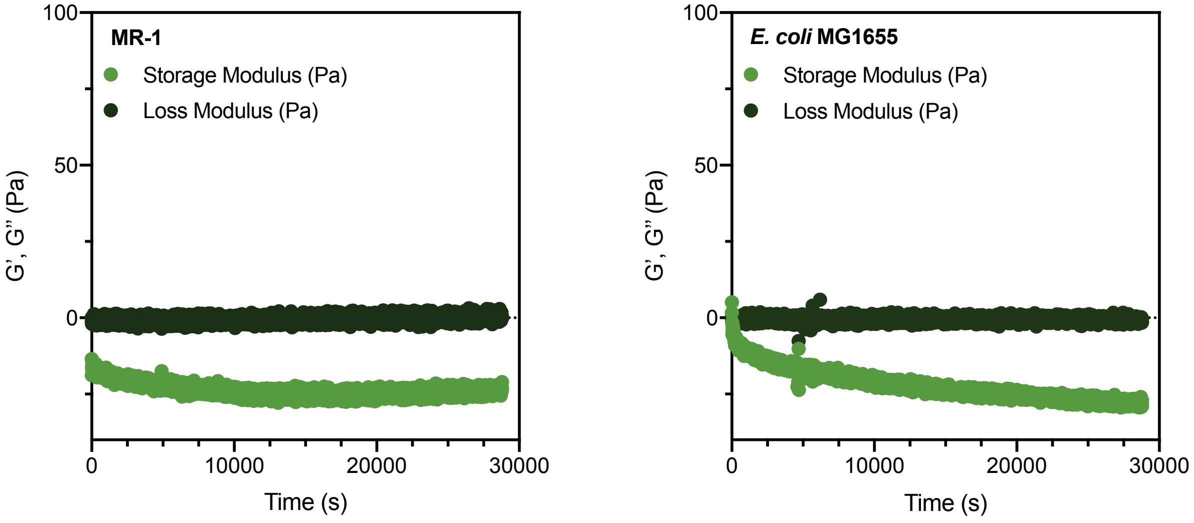


**Figure S5.** *In situ* rheology showing G' and G'' of non-functionalized HA solutions containing all crosslinking components and either MR-1 (left) or *E. coli* MG1655 (right). Gels did not form; negative G' is attributed to dominant inertial forces and small plate geometry (weak elastic component) (3).

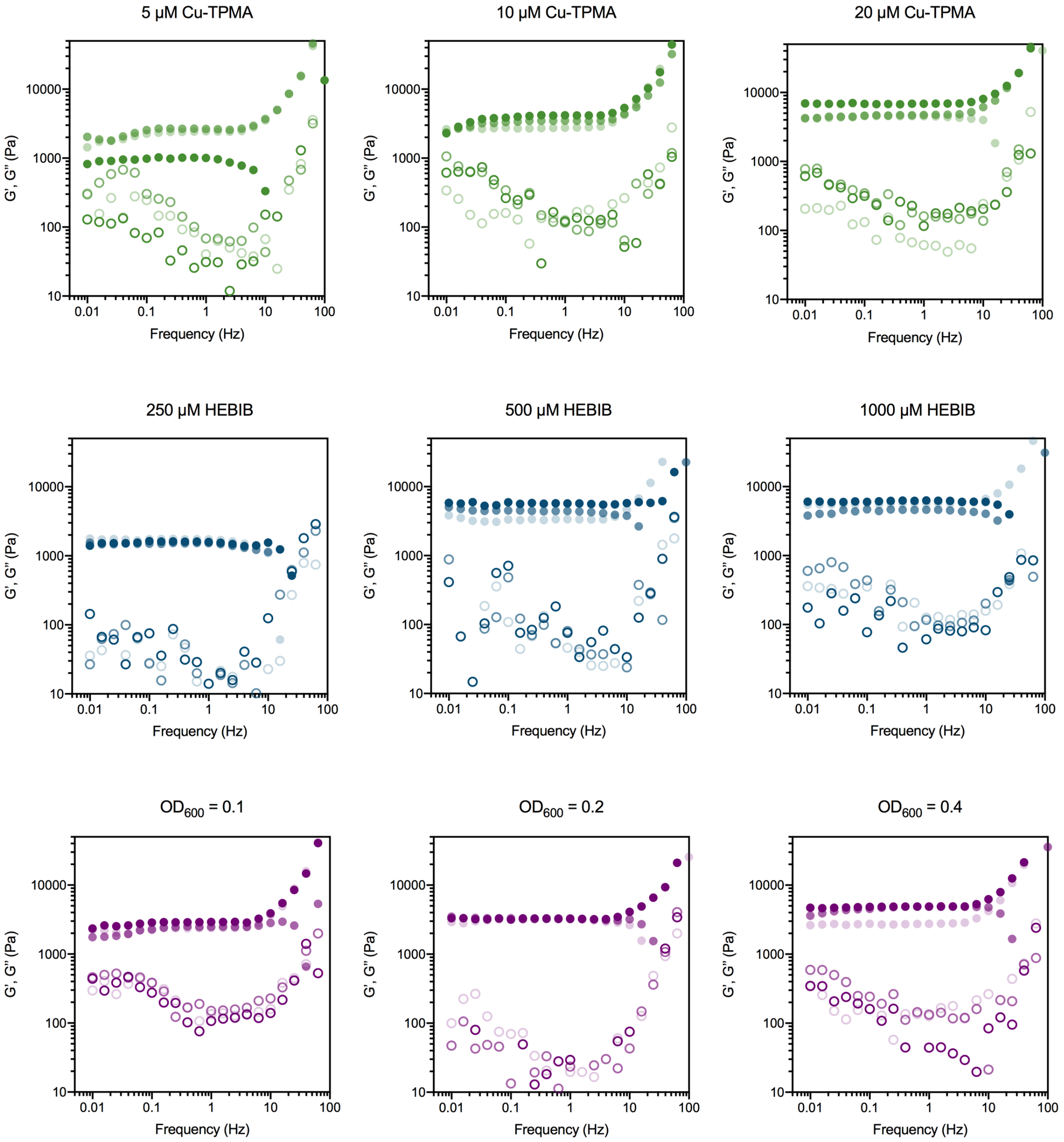


**Figure S6.** Rheological frequency sweeps for end-point measurements in Fig. 2. G' (filled circles) and G'' (open circles) for individual replicates.

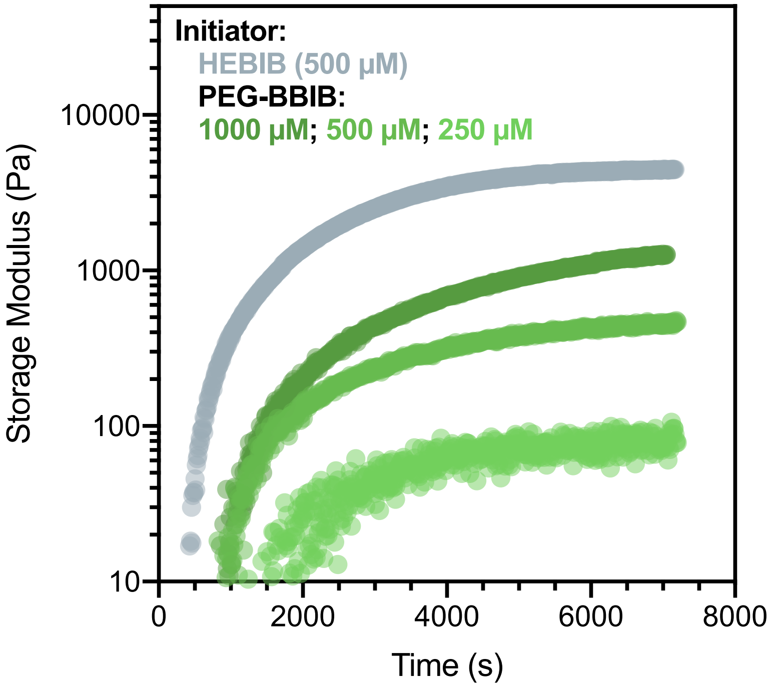


**Figure S7.** *In situ* G' measurement of MeHA65 gels synthesized using varying concentrations of poly(ethylene glycol) bis(2-bromoisobutyrate) ($M_{n,avg}$ = 700 g/mol, PEGBBiB). HEBiB, the initiator for standard gelation conditions, is included for comparison.

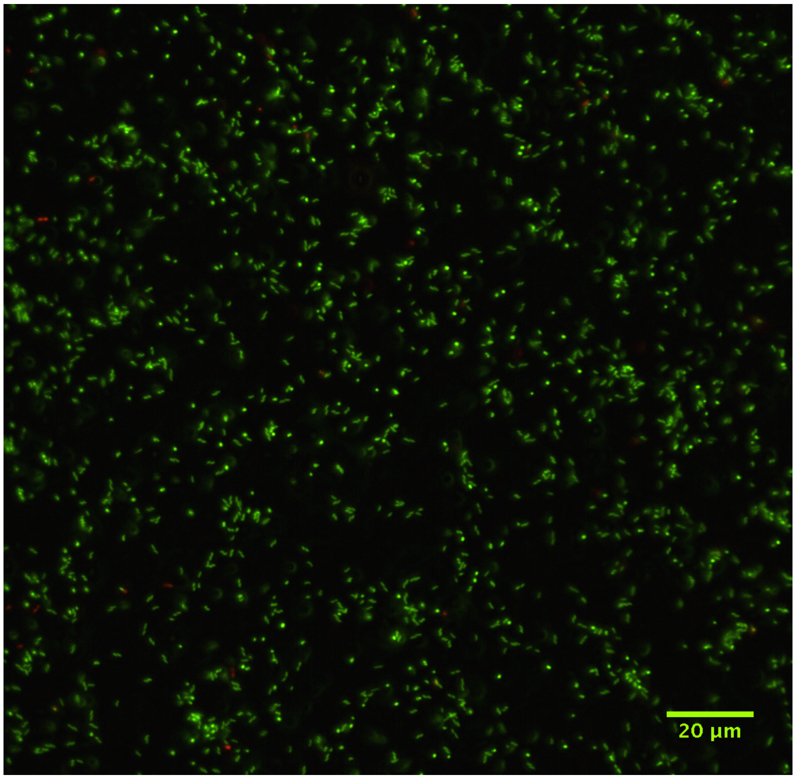


**Figure S9.** Viability analysis 5 days after crosslinking as determined by the BacLight Live/Dead stain. Cells are almost completely viable (green) as opposed to non-viable (red).

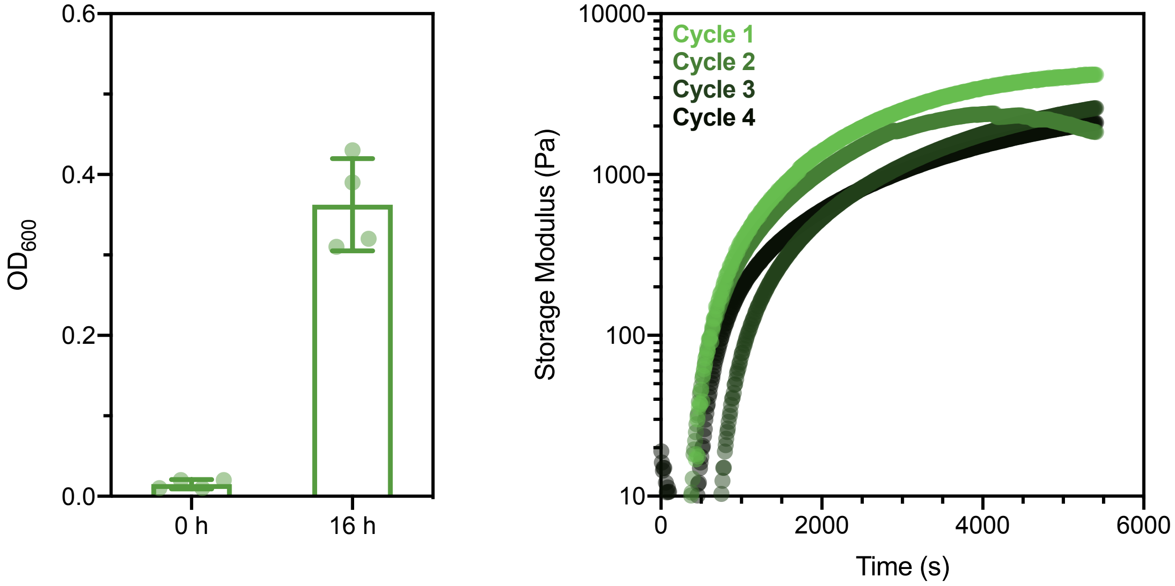


**Figure S10.** Cells can be regenerated after crosslinking to synthesize new gels. Left, $OD_{600}$ measurements of cultures after inoculating from swelling mixture into growth media, data are shown as mean ± SEM of $n$ = 4 measurements. Right, *in situ* storage modulus measurements (right) of gels crosslinked by recovered bacteria cultures (inoculated from hydrogel swelling media and grown overnight).

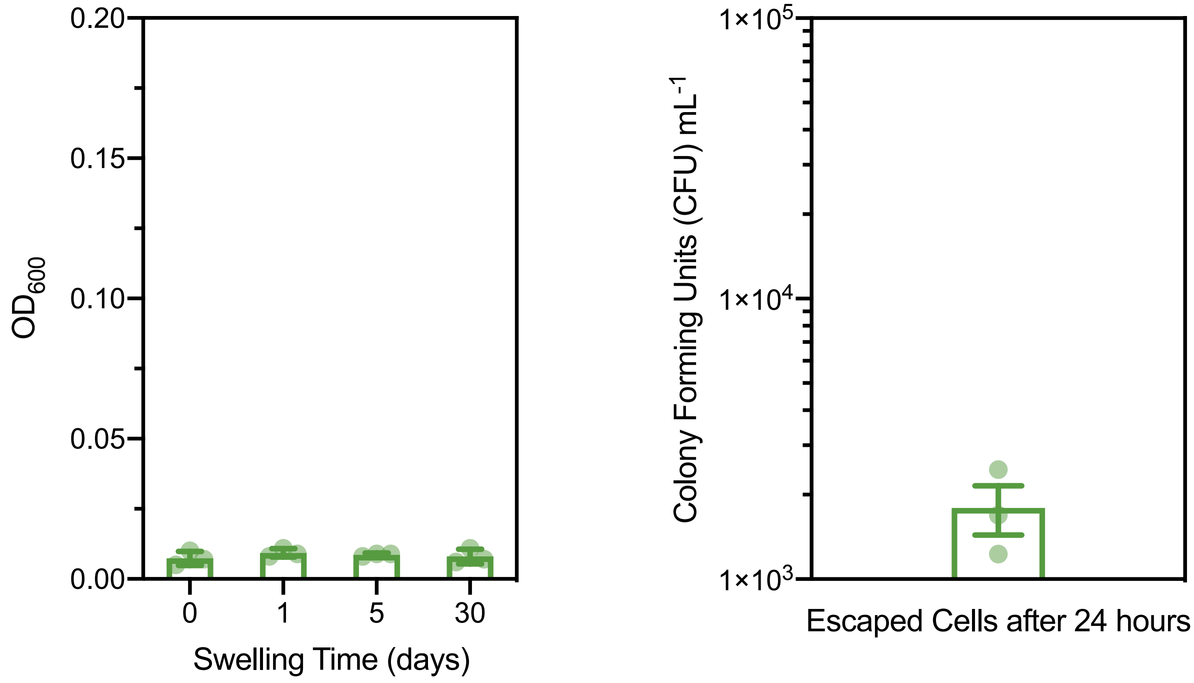


**Figure S11.** Bacteria do not appreciably escape the gels after synthesis. $OD_{600}$ measurements of swelling media (left) after washing gels 3x in PBS and allowing to swell in 1 mL PBS for varying lengths of time. Colony counting (right) confirms that escaped cells account for < 0.005% of inoculating density (~4 x $10^7$ CFU/mL or $OD_{600}$ = 0.2) after 3x washing in PBS. Data are shown as mean ± SEM for $n$ = 3 biological replicates.

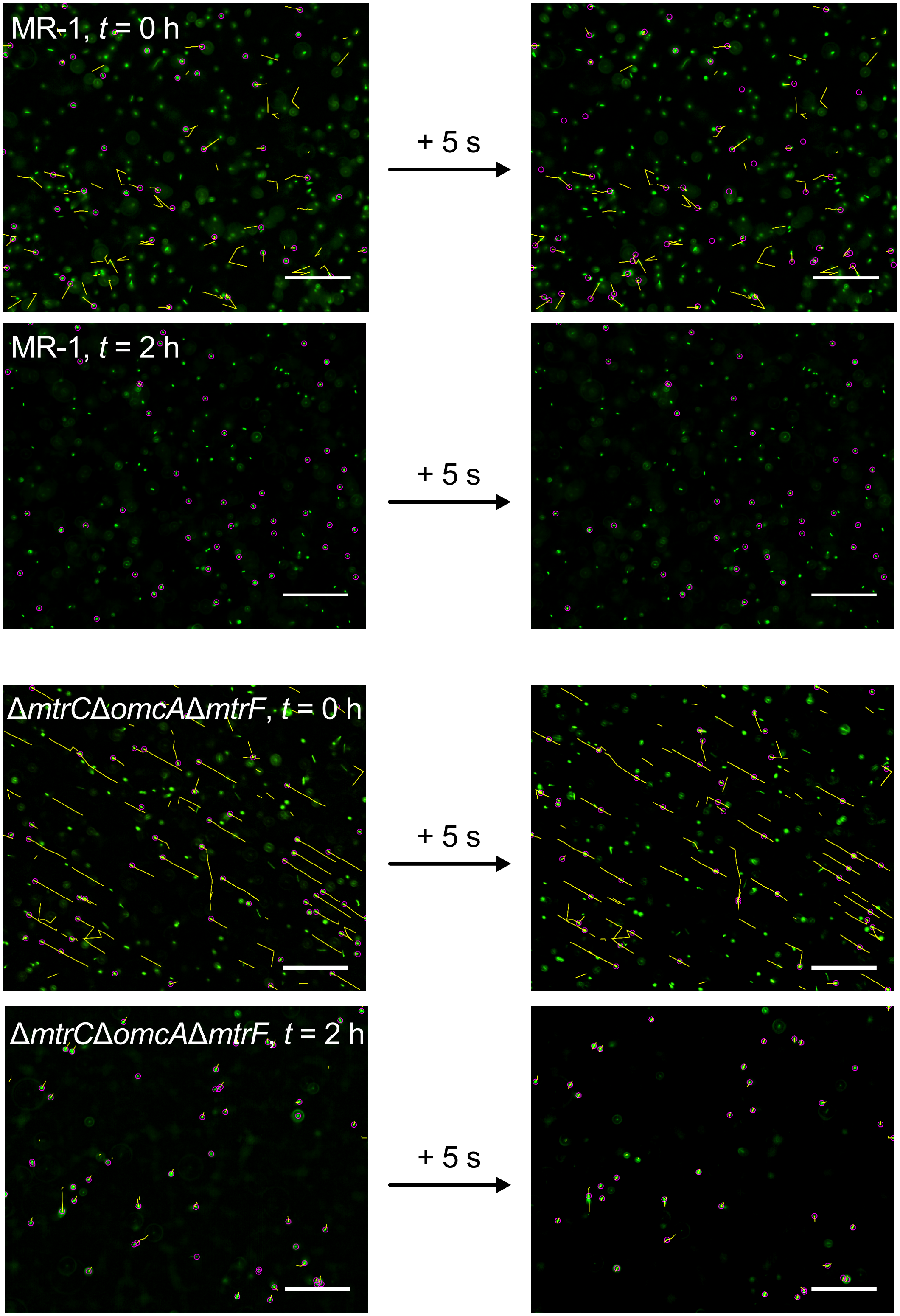


**Figure S12.** Fluorescence microscopy images of 1000 μM IPTG induced MR-1 and Δ*mtrC*Δ*omcA*Δ*mtrF* strains show swimming and convective movement over 5 seconds. Cells (purple circles) were tracked over 5 s (yellow lines) using TrackMate in Fiji 1.0. The average displacement of highest quality tracks (as determined by the software, *n* = 33) was then calculated for each strain at 0 and 2 hours.

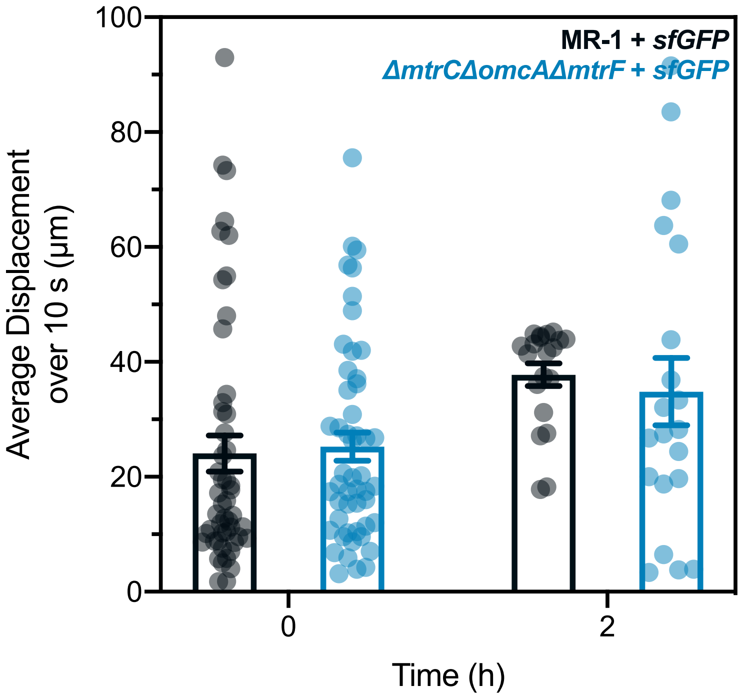


**Figure S13.** Average cell displacement within a non-functionalized 3 wt.% HA solution as measured by microscopy over 10 s time-lapses at 0 and 2 hours. Displacement was quantified using TrackMate in Fiji 1.0. Student t-test shows no statistical difference ($p$ = 0.95 at 0 hours, $n$ = 50; $p$ = 0.76 at 2 hours, $n$ = 20), indicating that differences in the functionalized MeHA is due to crosslinking. Data are shown as mean ± SEM.

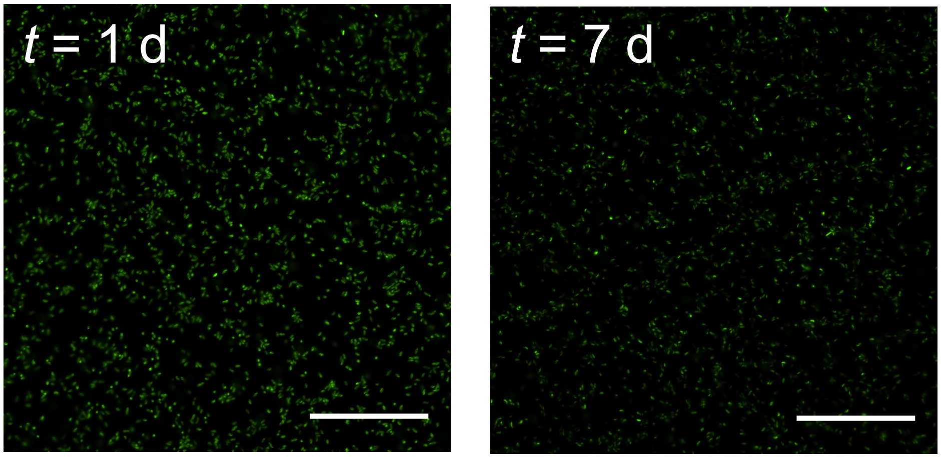


**Figure S14.** sfGFP fluorescence demonstrates metabolic activity for extended time periods after crosslinking. Crosslinked gels were swollen in 1x PBS for 1 and 7 days after crosslinking, then moved into 1000 µM IPTG-containing PBS for 24 hours and imaged by microscopy. Gels imaged before overnight induction with IPTG did not fluoresce. All images taken with equivalent exposure times to allow comparison of protein expression and circuit maintenance.

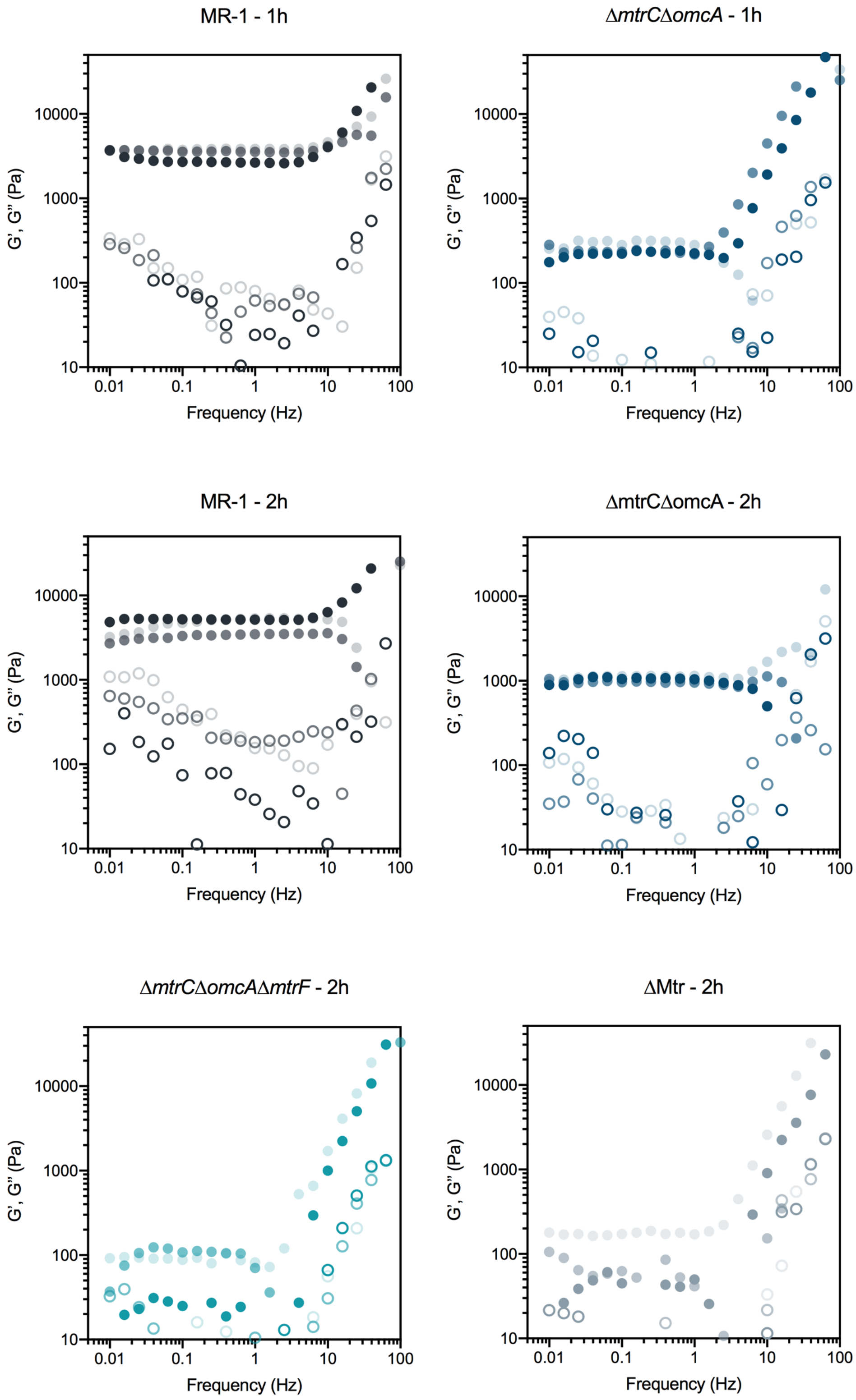


**Figure S15.** Rheological frequency sweeps for end-point measurements in Fig. 3. G' (filled circles) and G'' (open circles) for individual replicates.

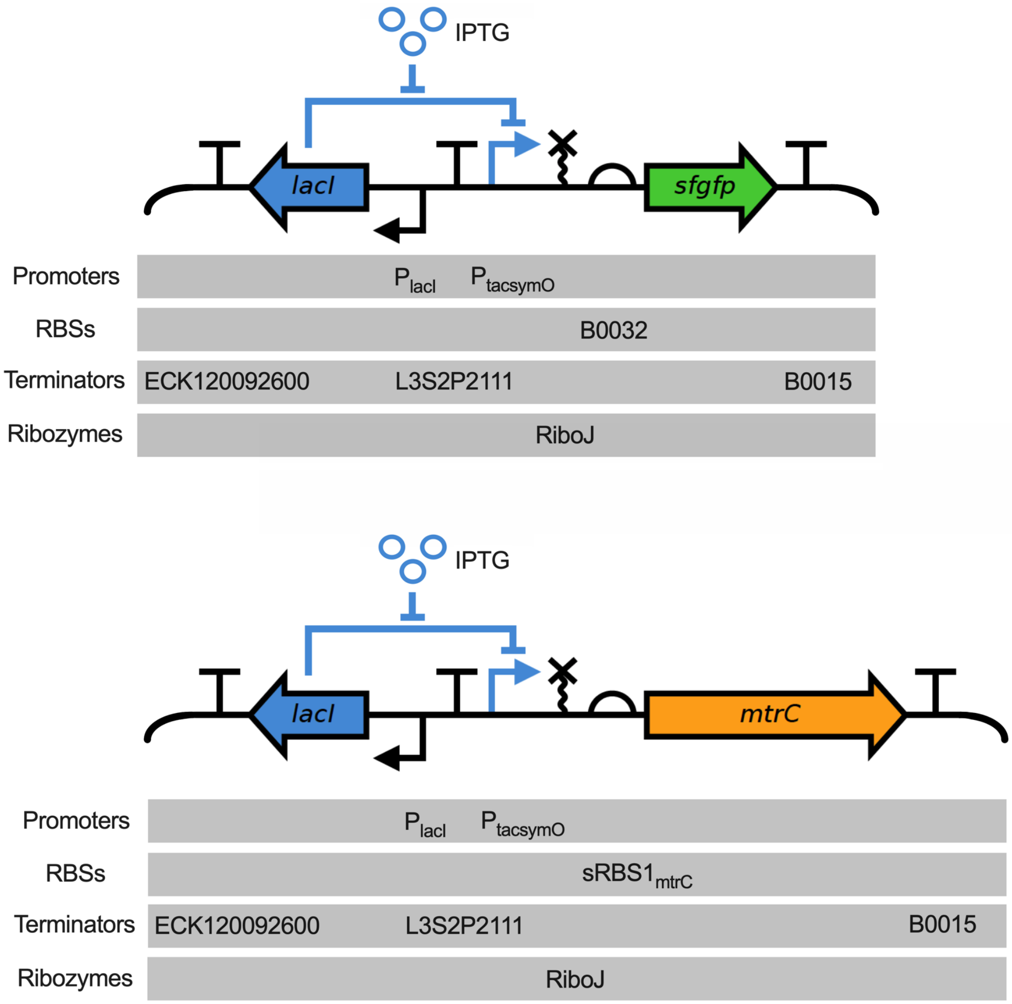


**Figure S16.** Gene circuit maps for both the *sfgfp* (top, pCD7sfGFP) and *mtrC* (bottom, pCD24r1) expression vectors. Outside of the flanking terminators, both vectors are identical and contain the ColE1 origin of replication and a kanamycin resistance marker. The empty vector (pCD8) is identical to pCD7sfGFP, but lacks *sfgfp*.

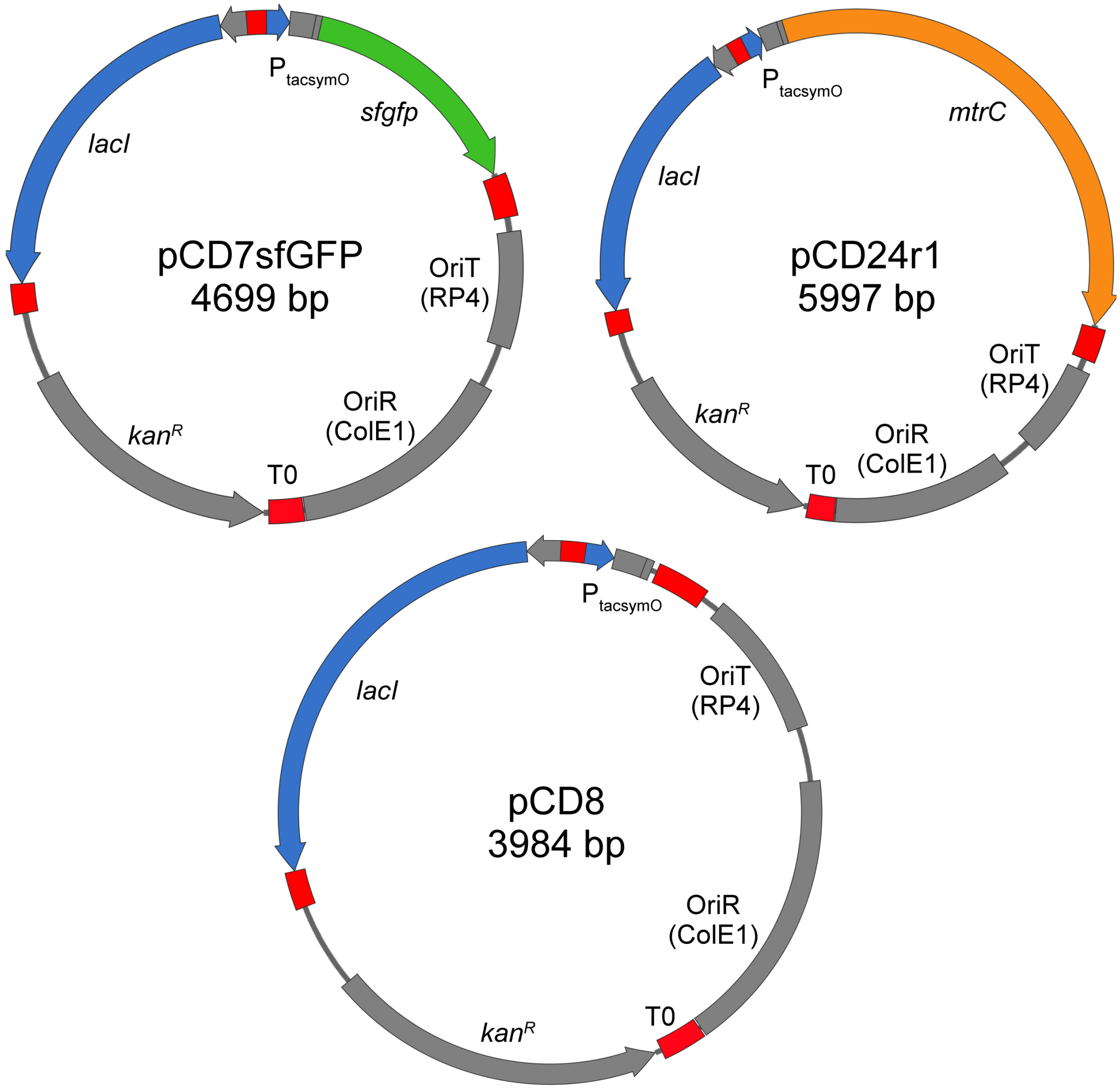


**Figure S17.** Plasmid maps for all plasmids used in this study. Terminator sequences are depicted as red, *lacI* and $P_{tacsymO}$ as blue, *sfgfp* as green, and *mtrC* as orange.

**Table S4.** Genetic parts/sequences used to construct the plasmids in this study.

| Genetic Part | DNA Sequence (5' to 3') |
|---|---|
| **Promoters** | |
| $P_{tacsymO}$ | TGTTGACAATTAATCATCGGCTCGTATAATGTGTGGAATTGTGAGCGCTCACAATTCTATGGACTATGTTT |
| $P_{lacI}$ | GCGGCGCGCCATCGAATGGCGCAAAACCTTTCGCGGTATGGCATGATAGCGCCCGGAAGAGAGTCAATTCAGGGTGGTGAAT |
| **Ribosome Binding Sites** | |
| B0032 | TCACACAGGAAAGTACTAG |
| $sRBS1_{mtrC}$ | GGGGAAAAACAGCAGTGCGAT |
| **Terminators** | |
| ECK120029600 | TTCAGCCAAAAAACTTAAGACCGCCGGTCTTGTCCACTACCTTGCAGTAATGCGGTGGACAGGATCGGCGGTTTTCTTTTCTCTTCTCAA |
| L3S2P2111 | TCGGTACCAAATTCCAGAAAAGAGGCCTCCCGAAAGGGGGGCCTTTTTTCGTTTTGGTCC |
| B0015 | TAATCTAGACCAGGCATCAAATAAAACGAAAGGCTCAGTCGAAAGACTGGGCCTTTCGTTTTATCTGTTGTTTGTCGGTGAACGCTCTCTACTAGAGTCACACTGGCTCACCTTCGGGTGGGCCTTTCTGCGTTTATA |
| **Ribozymes** | |
| RiboJ | AGCTGTCACCGGATGTGCTTTCCGGTCTGATGAGTCCGTGAGGACGAAACAGCCTCTACAAATAATTTTGTTTAA |
| **Genes** | |
| *lacI* | GTGAAACCAGTAACGTTATACGATGTCGCAGAGTATGCCGGTGTCTCTTATCAGACCGTTTCCCGCGTGGTGAACCAGGCCAGCCACGTTTCTGCGAAAACGCGGGAAAAAGTGGAAGCGGCGATGGCGGAGCTGAATTACATTCCCAACCGCGTGGCACAACAACTGGCGGGCAAACAGTCGTTGCTGATTGGCGTTGCCACCTCCAGTCTGGCCCTGCACGCGCCGTCGCAAATTGTCGCGGCGATTAAATCTCGCGCCGATCAACTGGGTGCCAGCGTGGTGGTGTCGATGGTAGAACGAAGCGGCGTCGAAGCCTGTAAAGCGGCGGTGCACAATCTTCTCGCGCAACGCGTCAGTGGGCTGATCATTAACTATCCGCTGGATGACCAGGATGCCATTGCTGTGGAAGCTGCCTGCACTAATGTTCCGGCGTTATTTCTTGATGTCTCTGACCAGACACCCATCAACAGTATTATTTTCTCCCATGAAGACGGTACGCGACTGGGCGTGGAGCATCTGGTCGCATTGGGTCACCAGCAAATCGCGCTGTTAGCGGGCCCATTAAGTTCTGTCTCGGCGCGTCTGCGTCTGGCTGGCTGGCATAAATATCTCACTCGCAATCAAATTCAGCCGATAGCGGAACGGGAAGGCGACTGGAGTGCCATGTCCGGTTTTCAACAAACCATGCAAATGCTGAATGAGGGTATCGTTCCCACTGCGATGCTGGTTGCCAACGATCAGATGGCGCTGGGCGCAATGCGCGCCATTACCGAGTCCGGGCTGCGCGTTGGTGCGGATATCTCGGTAGTGGGATACGACGATACCGAAGACAGCTCATGTTATATCCCGCCGTTAACCACCATCAAACAGGATTTTCGCCTGCTGGGGCAAACCAGCGTGGACCGCTTGCTGCAACTCTCTCAGGGCCAGGCGGTGAAGGGCAATCAGCTGTTGCCCGTGTCACTGGTGAAAAGAAAAACCACCCTGGCGCCCAATACGCAAACCGCCTCTCCCCGCGCGTTGGCCGATTCATTAATGCAGCTGGCACGACAGGTTTCCCGACTGGAAAGCGGGCAGTGA |
| *sfgfp* | ATGCGTAAAGGCGAAGAGCTGTTCACTGGTGTCGTCCCTATTCTGGTGGAACTGGATGGTGATGTCAACGGTCATAAGTTTTCCGTGCGTGGCGAGGGTGAAGGTGACGCAACTAATGGTAAACTGACGCTGAAGTTCATCTGTACTACTGGTAAACTGCCGGTACCTTGGCCGACTCTGGTAACGACGCTGACTTATGGTGTTCAGTGCTTTGCTCGTTATCCGGACCATATGAAGCAGCATGACTTCTTCAAGTCCGCCATGCCGGAAGGCTATGTGCAGGAACGCACGATTTCCTTTAAGGATGACGGCACGTACAAAACGCGTGCGGAAGTGAAATTTGAAGGCGATACCCTGGTAAACCGCATTGAGCTGAAAGGCATTGACTTTAAAGAAGACGGCAATATCCTGGGCCATAAGCTGGAATACAATTTTAACAGCCACAATGTTTACATCACCGCCGATAAACAAAAAAATGGCATTAAAGCGAATTTTAAAATTCGCCACAACGTGGAGGATGGCAGCGTGCAGCTGGCTGATCACTACCAGCAAAACACTCCAATCGGTGATGGTCCTGTTCTGCTGCCAGACAATCACTATCTGAGCACGCAAAGCGTTCTGTCTAAAGAT |

| | |
|---|---|
| | CCGAACGAGAAACGCGATCATATGGTTCTGCTGGAGTTCGTAACCGCAGCGG GCATCACGCATGGTATGGATGAACTGTACAAATGATGA |
| *mtrC* | ATGATGAACGCACAAAAATCAAAAATCGCACTGCTGCTCGCAGCAAGTGCCG TCACAATGGCCTTAACCGGCTGTGGTGGAAGCGATGGTAATAACGGCAATGA TGGTAGTGATGGTGGTGAGCCAGCAGGTAGCATCCAGACGTTAAACCTAGAT ATCACTAAAGTAAGCTATGAAAATGGTGCACCTATGGTCACTGTTTTCGCCAC TAACGAAGCCGACATGCCAGTGATTGGTCTCGCAAATTTAGAAATCAAAAAAG CACTGCAATTAATACCGGAAGGGGCGACAGGCCCAGGTAATAGCGCTAACTG GCAAGGCTTAGGCTCATCAAAGAGCTATGTCGATAATAAAAACGGTAGCTATA CCTTTAAATTCGACGCCTTCGATAGTAATAAGGTCTTTAATGCTCAATTAACGC AACGCTTTAACGTTGTTTCTGCTGCGGGTAAATTAGCAGACGGAACGACCGTT CCCGTTGCCGAAATGGTTGAAGATTTCGACGGCCAAGGTAATGCGCCGCAAT ATACAAAAAAATATCGTTAGCCACGAAGTATGTGCTTCTTGCCACGTAGAAGGT GAAAAGATTTATCACCAAGCTACTGAAGTCGAAACTTGTATTTCTTGCCACACT CAAGAGTTTGCGGATGGTCGCGGCAAACCCCATGTCGCCTTTAGTCACTTAA TTCACAATGTGCATAATGCCAACAAAGCTTGGGGCAAAGACAATAAAATCCCT ACAGTTGCACAAAATATTGTCCAAGATAATTGCCAAGTTTGTCACGTTGAATC CGACATGCTCACCGAGGCAAAAAACTGGTCACGTATTCCAACAATGGAAGTC TGTTCTAGCTGTCACGTAGACATCGATTTTGCTGCGGGTAAAGGCCACTCTCA ACAACTCGATAACTCCAACTGTATCGCCTGCCATAACAGCGACTGGACTGCT GAGTTACACACAGCCAAAACCACCGCAACTAAGAACTTGATTAATCAATACGG TATCGAGACTACCTCGACAATTAATACCGAAACTAAAGCAGCCACAATTAGTG TTCAAGTTGTAGATGCGAACGGTACTGCTGTTGATCTCAAGACCATCCTGCCT AAAGTGCAACGCTTAGAGATCATCACCAACGTTGGTCCTAATAATGCAACCTT AGGTTATAGTGGCAAAGATTCAATATTTGCAATCAAAAATGGAGCTCTTGATC CAAAAGCTACTATCAATGATGCTGGCAAACTGGTTTATACCACTACTAAAGAC CTCAAACTTGGCCAAAACGGCGCAGACAGCGACACAGCATTTAGCTTTGTAG GTTGGTCAATGTGTTCTAGCGAAGGTAAGTTTGTAGACTGTGCAGACCCTGC ATTTGATGGTGTTGATGTAACTAAGTATACCGGCATGAAAGCGGATTTAGCCT TTGCTACTTTGTCAGGTAAAGCACCAAGTACTCGCCACGTTGATTCTGTTAAC ATGACAGCCTGTGCCAATTGCCACACTGCTGAGTTCGAAATTCACAAAGGCA AACAACATGCAGGCTTTGTGATGACAGAGCAACTATCACACACCCAAGATGCT AACGGTAAAGCGATTGTAGGCCTTGACGCATGTGTGACTTGTCATACTCCTGA TGGCACCTATAGCTTTGCCAACCGTGGTGCGCTAGAGCTAAAACTACACAAA AAACACGTTGAAGATGCCTACGGCCTCATTGGTGGCAATTGTGCCTCTTGTCA CTCAGACTTCAACCTTGAGTCTTTCAAGAAGAAAGGCGCATTGAATACTGCCG CTGCAGCAGATAAAACAGGTCTATATTCTACGCCGATCACTGCAACTTGTACT ACCTGTCACACAGTTGGCAGCCAGTACATGGTCCATACGAAAGAAACCCTGG AGTCTTTCGGTGCAGTTGTTGATGGCACAAAAGATGATGCTACCAGTGCGGC ACAGTCAGAAACCTGTTTCTACTGCCATACCCCAACAGTTGCAGATCACACTA AAGTGAAAATGTAA |

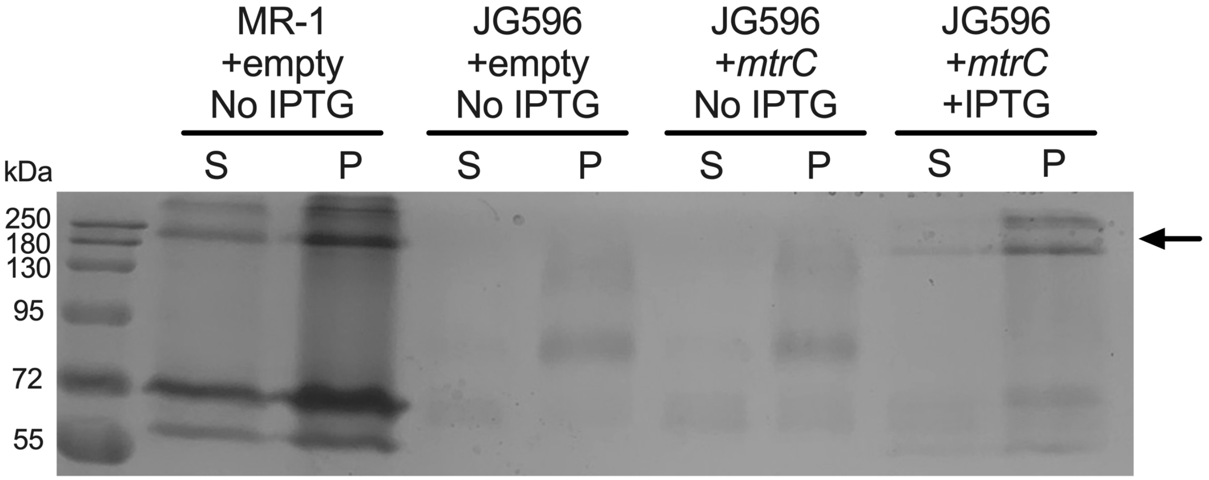


**Figure S18.** Heme-stained SDS-PAGE gel of uninduced (0 µM IPTG) and induced (1000 µM IPTG) *S. oneidensis* strains. 10 µg of protein (as determined by Bradford assay) was loaded into each lane. +empty indicates strains harboring the empty expression vector (pCD8) and +*mtrC* indicates strains harboring the IPTG-inducible *mtrC* vector (pCD24r1). After sonication, cell lysate was pelleted at 10,000 rcf for 5 minutes and the soluble fraction (S) and pellet (P) were separated for analysis. The arrow indicates the bands that appear in the JG596 +*mtrC* +IPTG lanes, which correspond to the apparent size of the MtrCAB complex (~210 kDa) (4).

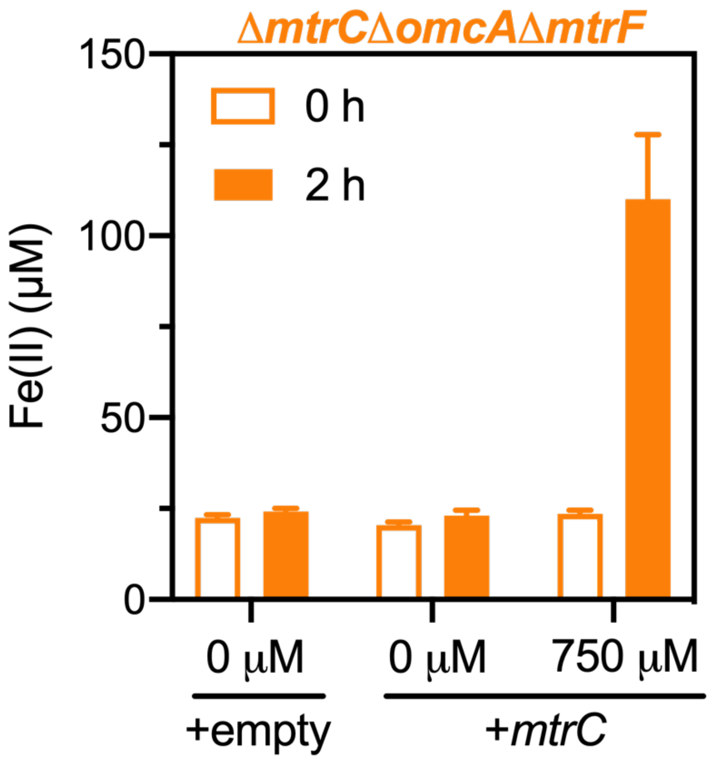


**Figure S19.** Reduction of Fe(III)-citrate over 2 hours by *S. oneidensis* Δ*mtrC*Δ*omcA*Δ*mtrF* carrying either pCD24r1 (+*mtrC*) or pCD8 (+empty). Cells were either uninduced (0 µM) or induced (750 µM) with IPTG at 0 h. Data shown are mean ± SD for *n* = 3 biological replicates.

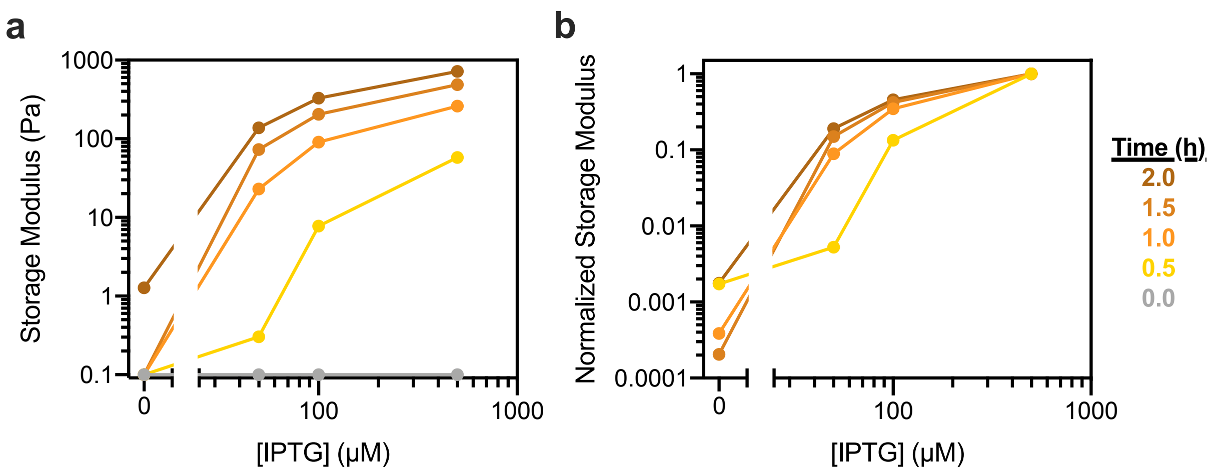


**Figure S20.** Dynamic range of storage modulus response function changes with time. (a) Storage moduli of hydrogels crosslinked by *S. oneidensis* Δ*mtrC*Δ*omcA*Δ*mtrF* carrying pCD24r1 (+*mtrC*) and (un)induced at various IPTG levels. Storage modulus was arbitrarily set at 0.1 Pa when stiffness was below the limit of detection. (b) Storage moduli from each timepoint when normalized to the highest IPTG concentration (500 μM). Past 1.0 h, all response curves collapse to similar functions. Each curve represents data from a single timepoint. Data shown are values from individual *in situ* rheology experiments (n = 1 biological replicate) that are also depicted in Figure 4b.

**Note S1.** Models of inducible gene expression and storage moduli.

**Hill Function Model**

The activating Hill function has been extensively used to describe inducible gene expression (5). Its general form is depicted below:

$$y = Bottom + (Top - Bottom)\frac{[I]^n}{{EC_{50}}^n + [I]^n} \quad [1]$$

where $y$ corresponds to storage modulus ($G'$) or relative expression units ($REU$), $Bottom$ corresponds to the lower-bound plateau value, $Top$ corresponds to the upper-bound plateau value, $[I]$ corresponds to IPTG concentration, $n$ corresponds to the hillslope, and $EC_{50}$ corresponds to the half-maximal effective concentration.

**Derivation of Linear Relationship Between REU and G'**

We observed that inducible transcriptional rate/sfGFP fluorescence ($REU$, Figure 4c) and hydrogel storage modulus ($G'$, Figure 4d) fit well to [1] and that similar $n$ and $EC_{50}$ values were obtained from fitting the model to each data set. Given this, we can substitute all $[I]$, $n$ and $EC_{50}$ terms with an expression containing $REU$ to obtain a linear relationship where $G' = f(REU)$. The final relationship is depicted below:

$$G' = \left(Bottom_{G'} - \frac{(Top_{G'} - Bottom_{G'})}{(Top_{REU} - Bottom_{REU})}Bottom_{REU}\right) + \frac{(Top_{G'} - Bottom_{G'})}{(Top_{REU} - Bottom_{REU})}REU \quad [2]$$

Assuming $[MtrC]$ is proportional to transcriptional rate ($REU$) (6), this generates a linear relationship between $G'$ and $[MtrC]$.

**Table S5.** Nonlinear fit parameters for products of *sfgfp* expression (fluorescence) and *mtrC* expression (storage modulus)

| Parameter | *sfgfp* Nonlinear Fit Value (units) | *mtrC* Nonlinear Fit Value (units) |
|---|---|---|
| **Best-fit Values** | | |
| Hillslope | 1.569 | 1.402 |
| $EC_{50}$ | 98.58 (μM) | 96.38 (μM) |
| Top | 1.032 (REU) | 3360 (Pa) |
| Bottom | 0.2086 (REU) | 158.4 (Pa) |
| **95% Confidence Interval** | | |
| Hillslopse | 1.028 to 2.505 | 0.3979 to 4.671 |
| $EC_{50}$ | 78.01 to 132.5 (μM) | 50.92 to 3650 (μM) |
| Top | 0.9654 to 1.137 (REU) | 2730 to 9795 (Pa) |
| Bottom | 0.1475 to 0.2644 (REU) | -401.7 to 568.6 (Pa) |
| **Goodness of Fit** | | |
| $R^2$ | 0.9314 | 0.8632 |
